# A novel non-genetic murine model of hyperglycemia and hyperlipidemia-associated aggravated atherosclerosis

**DOI:** 10.1101/2021.11.11.468191

**Authors:** Susanne Gaul, Khurrum Shahzad, Rebekka Medert, Ihsan Gadi, Christina Mäder, Dagmar Schumacher, Angela Wirth, Saira Ambreen, Sameen Fatima, Jes-Niels Boeckel, Hamzah Khawaja, Jan Haas, Maik Brune, Peter P Nawroth, Berend Isermann, Ulrich Laufs, Marc Freichel

## Abstract

**Objective:** Atherosclerosis, the main pathology underlying cardiovascular diseases is accelerated in diabetic patients. Genetic mouse models require breeding efforts which are time-consuming and costly. Our aim was to establish a new nongenetic model of inducible metabolic risk factors that mimics hyperlipidemia, hyperglycemia, or both and allows the detection of phenotypic differences dependent on the metabolic stressor(s).

**Methods and Results:** Wild-type mice were injected with gain-of-function PCSK9^D377Y^ (proprotein convertase subtilisin/kexin type 9) mutant adeno-associated viral particles (AAV) and streptozotocin and fed either a high-fat diet (HFD) for 12 or 20 weeks or a high-cholesterol/high-fat diet (Paigen diet, PD) for 8 weeks. To evaluate atherosclerosis, two different vascular sites (aortic sinus and the truncus of the brachiocephalic artery) were examined in the mice. Combined hyperlipidemic and hyperglycemic (HGHCi) mice fed a HFD or PD displayed characteristic features of aggravated atherosclerosis when compared to hyperlipidemia (HCi HFD or PD) mice alone. Atherosclerotic plaques of HGHCi HFD animals were larger, showed a less stable phenotype (measured by the increased necrotic core area, reduced fibrous cap thickness, and less α-SMA-positive area) and had more inflammation (increased plasma IL-1β level, aortic pro-inflammatory gene expression, and MOMA-2-positive cells in the BCA) after 20 weeks of HFD. Differences between the HGHCi and HCi HFD models were confirmed using RNA-seq analysis of aortic tissue, revealing that significantly more genes were dysregulated in mice with combined hyperlipidemia and hyperglycemia than in the hyperlipidemia-only group. The HGHCi-associated genes were related to pathways regulating inflammation (increased *Cd68, iNos*, and *Tnfa* expression) and extracellular matrix degradation (*Adamts4* and *Mmp14*). When comparing HFD with PD, the PD aggravated atherosclerosis to a greater extent in mice and showed plaque formation after 8 weeks. Hyperlipidemic and hyperglycemic mice fed a PD (HGHCi PD) showed less collagen (Sirius red) and increased inflammation (CD68-positive cells) within aortic plaques than hyperlipidemic mice (HCi PD). HGHCi-PD mice represent a directly inducible hyperglycemic atherosclerosis model compared with HFD-fed mice, in which atherosclerosis is severe by 8 weeks.

**Conclusion:** We established a nongenetically inducible mouse model allowing comparative analyses of atherosclerosis in HCi and HGHCi conditions and its modification by diet, allowing analyses of multiple metabolic hits in mice.

## Introduction

Atherosclerotic cardiovascular diseases, including coronary artery disease (CAD) and its complications, are a leading cause of mortality and morbidity globally (1). The risk of CAD is markedly increased in patients with both type 1 and type 2 diabetes mellitus (DM), with CAD events occurring earlier in patients with diabetes mellitus (2). Approximately 10% of total vascular deaths have been attributed to diabetes mellitus and its complications (2). Despite recent studies showing cardioprotective effects of new antidiabetic agents (3), there is a high need to understand how diabetes-associated alterations, particularly the evoked chronic hyperglycemia and metabolic alterations, aggravate atherosclerosis in CAD patients as a basis for the development of more effective treatments for these high-risk patients.

Currently used mouse models of hyperglycemia-associated atherosclerosis require a combination of streptozotocin (STZ) and crossbreeding with either apolipoprotein E (ApoE^-/-^) or low-density lipoprotein receptor (LDLR^-/-^) genetic KO mice, or alternatively, a double crossbreeding of ApoE^-/-^ or LDLR^-/-^ with insulin receptor (Ins2^+^, Akita) mutant mice (4). Some murine models, such as STZ-injected human apoB-expressing transgenic mice (5) or nonobese diabetic (NOD) mice (6), are resistant to the development of atherosclerosis. In addition, crossing mouse lines is time-consuming and costly. In particular, if mouse models harboring other genetic manipulations are to be used, backcrossing becomes a major issue. Thus, there is a high demand to generate an inducible mouse model of multiple metabolic hits, e.g., hyperlipidemia and hyperglycemia, which can be used alone or in combination with other genetic modifications. Bjorklund et al. developed a nongenetic mouse model of atherosclerosis induced by a single injection of recombinant adeno-associated virus (rAAV) encoding a hyperactive proprotein convertase subtilisin/kexin type 9 (PCSK9)^D377Y^ mutant followed by high-fat diet (HFD) feeding that has been used in nondiabetic settings (7, 8) as well as in mature diabetic Akita mice, where they showed diabetic-associated accelerated atherosclerosis (7). Here, we combined injections of rAAV8-PCSK9^D377Y^ and STZ with HFD feeding to generate a rapid and versatile method to induce hyperglycemia-induced aggravation of atherosclerosis in mice.

## Materials and Methods

All information regarding the materials and reagents is listed in the **Supplementary Major Resources Table**.

### Reagents

The following antibodies were used in the current study: rabbit anti-LDLR (R&D Systems, United States); mouse anti-β-actin (Abcepta Inc. United States); goat anti-rabbit IgG HRP (Cell Signaling Technology, Germany); rat anti-MOMA-2, rabbit anti-CD68 and rabbit anti-alpha smooth muscle actin (α-SMA) (Abcam, Germany); and rabbit anti-GAPDH (Sigma–Aldrich, Germany). The following secondary antibodies for immunofluorescence were used: Texas red rabbit anti-mouse IgG (Vector Laboratories, United States) and goat anti-rat IgG (H+L) cross-adsorbed secondary antibody, Alexa Fluor 568 (Thermo Fisher, United States). Other reagents were as follows: BCA reagent (Perbio Science, Germany); vectashield mounting medium with DAPI (Vector Laboratories, United States); nitrocellulose membrane (Bio–Rad, USA) and immobilon™ western chemiluminescent HRP substrate (Merck, Millipore, United States); streptozotocin (Enzo Life Sciences, Germany); Oil Red O (Sigma–Aldrich, Germany); Accu-Chek test strips, Accu-Check glucometer, and protease inhibitor cocktail (Roche Diagnostics, Germany); albumin fraction V, hematoxylin Gill II, acrylamide, and agarose (Carl ROTH, Germany); aqueous mounting medium (ZYTOMED, Germany); “high-fat diet” (HFD) experimental food (Western-type diet containing 21% fat and 0.21% cholesterol or Paigen diet containing 16% fat, 1.25% cholesterol (TD88137, Ssniff, Germany), and 0.5% sodium cholate, (D12336, Ssniff, Germany); PBS (Life Technologies, Germany); rompun 2% (Bayer, Germany); and ketamine 10% (beta-pharm, Germany).

### Mice

Eight-week-old male LDLR^-/-^ (002207) mice were obtained from the Jackson Laboratory (Bar Harbor, ME, USA). Male wild-type C57BL/6N mice were obtained from Charles River Laboratories (Wilmington, MA, USA). Only age-matched male mice were used throughout the study (6-7 mice per group). All animal experiments complied with the ARRIVE guidelines and were carried out in accordance with the Directive 2010/63/EU guidelines. They were conducted following standards and procedures approved by the local Animal Care and Use Committee (35-9185.81/G-185/19, Regierungspräsidium Karlsruhe, Germany).

### Generation and quantification of rAAV8 viral particles

Recombinant rAAV8 vector particles were generated and purified using the iodixanol gradient ultracentrifugation method(9, 10). rAAV8 production was carried out using HEK293T cells. First, 1.8 x 10^8^ HEK293T cells were seeded in a ten-chamber CellStack (Corning, USA) and cultured in DMEM+ Glutamax (Gibco, Thermo Fisher Scientific, USA) supplemented with 10% fetal bovine serum (FBS) and 1% penicillin G/streptomycin. After 48 hours, a 1:1:1 molar ratio of pAAV-D377YmPCSK9-bGHpA plasmid(7), the rep-cap AAV8 helper plasmid and an adenoviral helper plasmid was mixed and transfected using polyethylenimine (PEI) (Polyscience, USA). The cells were harvested in 3 ml of lysis buffer and lysed by four freeze–thaw cycles 72 hours after transfection. The vector particles were purified using an iodixanol (Progen, Germany) gradient consisting of four phases with decreasing density (60%, 40%, 25%, and 15%) and ultracentrifugation at 50,000 x g for 135 min at 4 °C. Approximately 3 ml of the 40% phase, in which predominantly full virus particles accumulated, was recovered with a 27G needle. Finally, the vector solution was buffered into PBS using dialysis tubes (Zeba Spin Desalting Columns 7K MWCO, Thermo Scientific, USA) and concentrated (VivaSpin 10K MWCO, Sartorius, Germany). The vector titer was quantified as genome copy numbers per milliliter using a qPCR SYBR-Green assay (Bio–Rad) and primer sequences specific for the bGHpA sequence (bGHpA-fw 5’ACCTAACTCACTGATCCGAAATTA 3’, bGHpA-rev 5’ATTTCGGATCAGTGAGTTAGG 3’)(11).

### Induction of atherosclerosis by hypercholesterolemia (HCi) and hypercholesterolemia and hyperglycemia (HGHCi) in mice

Adeno-associated viral vectors encoding the gain-of-function variant D377Y of murine PCSK9 (rAAV8-PCSK9^D377Y^) under the control of a liver-specific promoter were delivered via a single retro-orbital sinus injection (1.0 x 10^11^ viral genomes/mouse), and treated animals were fed either a high-fat Western-type diet (HFD, containing 21% fat and 0.21% cholesterol) or a high-cholesterol/high-fat Paigen diet (PD, containing 16% fat, 1.25% cholesterol, and 0.5% sodium cholate) to induce chronic hypercholesterolemia (HCi). Control animals not treated with rAAV8-PCSK9^D377Y^ were fed a high-fat diet (graphic abstract).

To induce chronic hyperglycemia and hypercholesterolemia (HGHCi), mice were injected with streptozotocin (STZ, 60 mg/kg, intraperitoneally, once daily for five consecutive days, freshly dissolved in 0.05 M sterile sodium citrate, pH 4.5) one week after rAAV8-PCSK9^D377Y^ application. As a control for STZ injections, mice received injections with an equal volume of 0.05 M sodium citrate, pH 4.5, for 5 days. As PCSK9 expression leads to the degradation of low-density lipoprotein (LDL) receptors, we quantitatively compared the development of atherosclerotic plaque formation in the inducible HCi and HGHCi models with that in LDLR^-/-^ mice fed a HFD or PD. LDLR^-/-^ mice served as the established control model for hypercholesterolemia-evoked atherosclerosis(12). Blood glucose levels were monitored twice a week using the Accu-Chek Aviva system (Roche, USA) and maintained in the range of 300–500 mg/dl. Body weight was measured once weekly. HFD and/or hyperglycemia (minimum 300 mg/dl) was maintained for up to 12 or 20 weeks. In the first 4 weeks of the study, some mice in the PD and HFD groups did not tolerate the food. These animals showed a strong reduction in body weight, which was defined as a termination criterion. Consequently, these mice were removed from the study. Mice fed the PD were analyzed after 8-9 weeks due to early mortality in the hyperglycemic group (**Supplementary Figure I**). At the respective study endpoints, mice were sacrificed, and atherosclerotic plaque morphology was analyzed as previously described(13–15).

### Analysis of mice

At the end of the study period (12 or 20 weeks for HFD or 8 weeks for PD), body weight was measured, and the mice were sacrificed(14–16). Blood samples were obtained from the inferior vena cava of anticoagulated mice (500 U of unfractionated heparin, intraperitoneally). Blood was centrifuged at 2,000 x g for 20 min at 4 °C, and plasma was snap frozen in liquid nitrogen. Mice were perfused with ice-cold PBS for 10 min, and the heart and aortic arches, including the brachiocephalic arteries, were embedded in O.C.T. compound and snap frozen. Brachiocephalic arteries (from distal to proximal) and upper hearts (aortic sinus) were sectioned at 5- and 10-μm thickness, respectively.

### Plasma analysis

Heparin plasma was prepared for the measurement of plasma lipids. A total of 500–700 μl of blood per mouse was collected in heparin tubes and centrifuged at 3,500 x g for 10 min at room temperature. Plasma was transferred to a 1.5-ml tube and stored at −80 °C. Plasma samples of HFD- and PD-fed mice were diluted 1:5 with 0.9% NaCl before cholesterol and triglyceride measurements. Plasma samples were analyzed in the accredited central laboratory of Heidelberg University Hospital using standard operating procedures according to the manufacturers’ instructions. Cholesterol and triglycerides were analyzed on a Siemens ADVIA Chemistry XPT System (reagent kits 04993681 and 10697575, respectively). We measured the concentrations of mouse IL-1β by ELISA (R&D Systems) according to the manufacturer’s instructions.

### Histology

Oil Red O staining was conducted on frozen cross-sections of the aortic sinus and the truncus of the brachiocephalic artery (BCA) (14, 17). Cryopreserved cross-sections of the brachiocephalic arteries and aortic sinus (5 and 10 μm, respectively) were fixed in ice-cold acetone for 2 min, rinsed twice in ice-cold 1x PBS, and stained with Oil Red O for 10 min. Sections were rinsed twice with distilled water for 20 seconds and once in running tap water for 10 min. Sections were then counterstained with hematoxylin for 40 seconds, rinsed in tap water, and mounted with aqueous mounting medium. Movat staining was performed on frozen sections of the aortic sinus and brachiocephalic arteries. Frozen sections (5 μm) were fixed in Bouin’s solution at 50 °C for 10 min and stained with 5% sodium thiosulfate for 5 min, 1% alcian blue for 15 min, alkaline alcohol for 10 min, Movat’s Weigert’s solution for 20 min, crocein scarlet acid/fuchsin solution for 1 min, 5% phosphotungstic acid for 5 min and 1% acetic acid for 5 min. Between every staining step, the tissue sections were washed with tap water and distilled water. Sections were then covered with cytoseal mounting medium. Every 15^th^ section (~90 μm) of the brachiocephalic arteries and aortic sinus were analyzed to quantify the plaque area. The following parameters were determined. 1) The vessel lumen, where the vessel lumen is the area within the blood vessel, consisting of both the remaining open lumen and the plaque area, that does not include the vessel wall itself. 2) Total plaque size, where the size of the plaque comprises all parts of the atheroscleroma (fibrous cap, necrotic tissue, fibrous tissue, etc.) within the vessel lumen. 3) The necrotic core (as a percentage) is defined as the area stained blue upon Movat’s pentachrome staining and is presented as the percentage of the total plaque size. 4) The fibrous cap, which is the minimal thickness of the fibrous tissue overlaying the necrotic core. If multiple necrotic cores were present within one plaque, the thickness of all fibrous caps was determined, and the average was used for further analyses. 5) the EEL, IEL and media, where the area surrounded by the external elastic lamina (EEL) and the internal elastic lamina (IEL) were measured using bright field images of Oil Red O-stained BCA images as described previously (18) (**Supplementary Figure II**). The EEL and IEL were encircled (**Supplementary Figure II**), and their areas were measured using Image-Pro Plus software. The media area was calculated by subtracting the IEL area from the EEL area. The lumen area was calculated by subtracting the plaque area from the IEL area. Thickness was measured using ImageJ software using a free-hand tool (13, 14). Cryosections of PD-fed mice were used for picrosirius red staining (Sigma–Aldrich, Germany) according to the manufacturer’s instructions. Tissue sections were then stained with picrosirius red solution for 1 h at room temperature. The sections were washed 2 times in acidified water (5 ml of glacial acetic acid to 1 liter of water) and mounted. For histological analysis, images were captured with a Keyence BZ-X810 fluorescence microscope (aortic sinus) and an Olympus Bx43 microscope (brachiocephalic arteries). Image-Pro Plus software (version 6.0) and ImageJ software were used for image analysis(14, 15, 17).

### Immunohistochemistry and immunofluorescence staining

For immunohistochemistry and immunofluorescence staining, frozen sections of brachiocephalic arteries (BCA) and aortic roots were fixed in ice-cold acetone for 8 min, washed twice with ice-cold PBS and incubated in 2% BSA in PBST for 1 h. Sections were then incubated overnight at 4 °C with primary antibodies against α-SMA (1:250) and CD68 (1:1000). Sections incubated without primary antibodies were used as negative controls for background correction. After overnight incubation, the sections were washed three times with PBS followed by incubation with corresponding horseradish peroxidase (HRP)-labeled secondary antibodies. After washing, tissues were counterstained using 3,3’-diaminobenzidine (DAB)/hematoxylin. Primary antibody against MOMA-2 (1:100) was incubated overnight at 4 °C, followed by washing three times with PBS and incubation with fluorescently labeled corresponding secondary antibody. Sections incubated without secondary antibody were used as negative controls and for background correction. After washing, nuclear counterstaining was conducted using mounting medium with DAPI. Images were captured and analyzed using a Keyence BZ-X810 all-in-one fluorescence microscope and an Olympus Bx43 microscope (Olympus, Hamburg, Germany) using the same settings in the experimental and control groups. Analyses were performed by two independent blinded investigators. ImageJ software (Version 1.8.0) was used for image analysis.

### Immunoblotting

Proteins were isolated, and immunoblotting was performed as previously described(19). Mouse livers were weighed, and an adjusted volume of RIPA buffer containing protease inhibitor cocktail (100 μl/10 mg) was added. Tissue was homogenized mechanically using 20G and 25G needles followed by a 30-min incubation on ice with sequential vortexing. Samples were centrifuged at 12,000 x g for 20 min at 4 °C. The supernatant was transferred to a fresh tube, and the protein concentration was determined using the Pierce™ BCA Protein Assay Kit following the manufacturer’s instructions and a Varioskan Lux plate reader (Thermo Fisher Scientific, Waltham, MA, USA). Samples were separated using the Mini-PROTEAN® TGX™ Precast Gel 4-15% (Bio–Rad Laboratories, Inc., Hercules CA, USA) and transferred to a nitrocellulose membrane. Membranes were incubated with Invitrogen No-Stain solution for 10 min and blocked for 1 h in 5% low-fat milk dissolved in TBS-T. Incubation with primary anti-mLDLR antibody and anti-β-actin was performed overnight at 4 °C. Membranes were incubated with the corresponding secondary rabbit anti-goat immunoglobulin/HRP and goat anti-mouse immunoglobulin/HRP for 1 h at room temperature. Millipore Immobilon Classico Western HRP Substrate was applied to detect the signal using iBright 1500 (Thermo Fisher Scientific, Waltham, MA, USA). Densiometric analysis was performed using ImageJ software.

### RNA-seq, functional annotation and pathway analysis

RNA was extracted from aortic tissues (comprising the plaque and surrounding tissue) using an RNeasy mini kit (QIAGEN, Germany), and the RNA concentration was measured using a Nanodrop (2000C, Peq lab, Germany). The quality and integrity of RNA were controlled with an Agilent Technologies 2100 Bioanalyzer (Agilent Technologies, Waldbronn, Germany). Expression profiling was performed using RNA sequencing (RNA-seq). An RNA sequencing library was generated from 500 ng of total RNA using a Dynabeads® mRNA DIRECT(tm) Micro Purification Kit (Thermo Fisher) for mRNA purification followed by a NEBNext® Ultra(tm) II Directional RNA Library Prep Kit (New England BioLabs) according to the manufacturer’s protocols. The libraries were sequenced on an Illumina NovaSeq 6000 using a NovaSeq 6000 S2 Reagent Kit (100 cycles, paired-end run) with an average of 3 x 10^7^ reads per RNA sample. A quality report was generated by the FASTQC (version 0.11.8) tool for each FASTQ file. Before alignment to the reference genome, each sequence in the raw FASTQ files was trimmed on base call quality and sequencing adapter contamination using the Trim Galore! wrapper tool (version 0.4.4). Reads shorter than 20 bp were removed from the FASTQ file. Trimmed reads were aligned to the reference genome using the open source short read aligner STAR (version 2.5.2b, https://code.google.com/p/rna-star/) with settings according to the log file. Feature counts were determined using the R package Rsubread (version 1.32.4). Only genes showing counts greater than 5 at least two times across all samples were considered for further analysis (data cleansing). Gene annotation was performed using the R package bioMaRt (version 2.38.0). Before starting the statistical analysis steps, expression data were log2 transformed and normalized according to the 50^th^ percentile (quartile normalization using edgeR, version 3.24.3). Differential gene expression was calculated by the R package edgeR. Statistically significant DEGs (*p*<0.05 and FDR <0.05) were sorted and categorized after correcting for multiple hypothesis testing by the Benjamini– Hochberg method. The threshold to identify differentially expressed genes (DEGs) was set to a logFc value of ± 0.58 (IRI group), resulting in a 1.5-fold expression change. To identify genes differentially regulated by either hyperlipidemia (HCi) or combined hyperglycemia and hyperlipidemia (HGHCi), DEGs between the HFD (control) group and either group were identified based on a minimum logFc difference value of ±0.58. Heatmapper (http://www.heatmapper.ca/) was used to generate heatmaps of gene expression data. Genes shown in the heatmap were sorted based on DEGs in the control group (HFD without PCSK9 injections) based on their log Fc values. For representation purposes, no clustering method was applied, and the z score was used. Venny (version 2.1), an online interactive tool, was used for comparison and identification of overlapping DEGs between different groups. Gene ontology was performed using the online tool Database for Annotation, Visualization and Integrated Discovery (DAVID) version 6.8 bioinformatics package, and Benjamini Hochberg adjustment was applied to all enriched p values to control for multiple testing. All RNA-seq analyses were performed by an independent blinded investigator.

### qPCR analysis

RNA was isolated using an RNeasy Mini kit from Qiagen, and reverse transcription was performed with iScript (Bio–Rad) Mastermix. Primer sequences and probes are listed in the Major Resources Table in the Supplemental Material. qPCR analyses were performed with PowerUp SYBR Green Mastermix (Thermo Fisher). All samples were run in duplicate, and relative gene expression was converted using the 2-ΔΔCT method against the mean of two internal control housekeeping genes, namely, hypoxanthine-guanine phosphoribosyl transferase (*Hprt*) and β-2 Microglobulin (*B2m*) for mice. ΔΔCT= (CTexperiment gene – CTmean experiment housekeeping) - (CTcontrol gene – CTmean control housekeeping). The relative gene expression in the control HFD group was set at 1.

### Statistical analysis

Statistical analyses were performed with GraphPad Prism (version 7; GraphPad Software Inc., La Jolla, CA, USA). The significance level was set at p< 0.05 for all comparisons. The data are summarized as the mean ± standard error of the mean (SEM). Comparisons of two groups were analyzed with unpaired Student’s T test. Statistical analyses of more than two groups were performed with analysis of variance (ANOVA) and Sidak’s post hoc comparisons. Differences between groups of a single independent variable were determined using one-way ANOVA, and between two independent variables using two-way ANOVA. The Kolmogorov–Smirnov test or D’Agostino-Pearson normality test was used to determine whether the data were consistent with a Gaussian distribution.

## Results

### The combination of hyperglycemia and hyperlipidemia exacerbates atherosclerosis in PCSK9^D377Y^-expressing mice fed a high-fat diet

To induce hypercholesterolemia (HCi), 8-week-old C57BL/6N mice were administered a single dose of rAAV8-PCSK9^D377Y^ (1.0 x 10^11^ adeno-associated viral particles, intravenously) one week before feeding a high-fat diet (HFD, Western-type diet). Saline-injected mice fed a HFD served as a control for rAAV8-PCSK9^D377Y^ intervention (control HFD). Saline-injected mice with genetic deficiency of low-density lipoprotein receptor (LDLR knockout (KO)) on a HFD served as controls for rAAV8-PCSK9^D377Y^ treatment (**Figure 1A**). The mouse group in which hypercholesterolemia evoked by PCSK9 expression and HFD (HCi, single hit) was combined with induction of chronic hyperglycemia using STZ injection (double hit) is hereafter termed HGHCi (high glucose high cholesterol - inducible) throughout the manuscript. For quantification of atherosclerotic plaque formation, mice were euthanized 12 (early time point) or 20 (late time point) weeks after rAAV8-PCSK9^D377Y^ injection and the start of HFD feeding.

**Figure 1:**
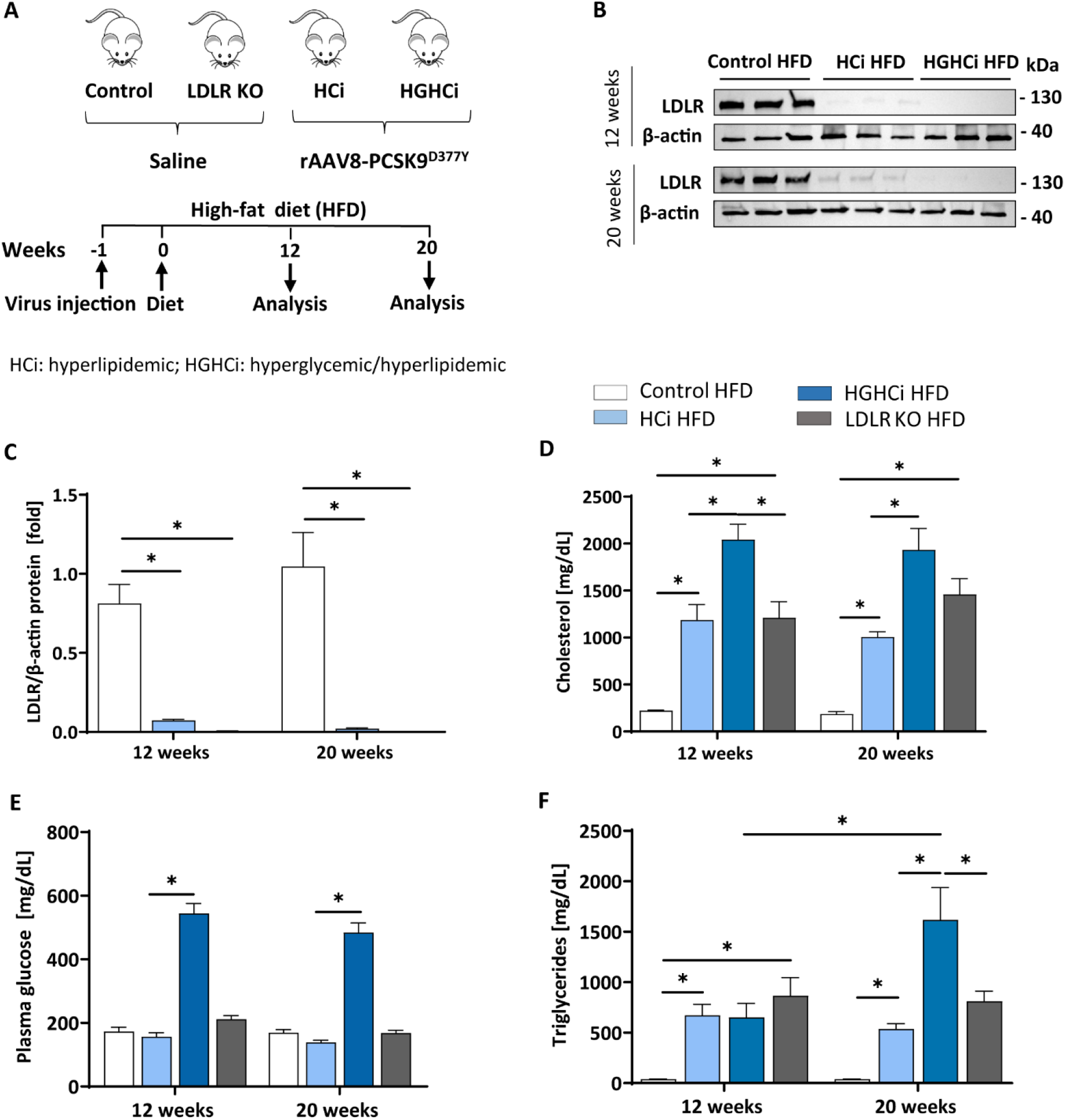
Simultaneous treatment of rAAV8-PCSK9 virus and Streptozotocin induces a hyperlipidemic and hyperglycemic mouse phenotype. **A:** Schematic summary of the experimental setup. Mice were analyzed after 12 or 20 weeks of interventions initiation. Wild type mice without rAAV8-PCSK9^D377Y^ injection on high-fat diet (HFD, 21 % fat, 0.21 % cholesterol) was used as control. Mice injected with rAAV8-PCSK9^D377Y^ and fed a HFD are termed HCi HFD (hyperlipidemic), or injected with both rAAV8-PCSK9^D377Y^ and streptozotocin and fed a HFD are termed HGHCi HFD (hyperlipidemic and hyperglycemic). LDLR KO mice on HFD served as atherosclerosis reference control. **B:** Representative immunoblot of liver lysate showing hepatic protein levels of low-density lipoprotein receptor (LDLR). β-actin was used as loading control. **C:** Densitometric analysis of LDLR immunoblot was normalized on β-actin and referred to control HFD group which is set at 1. Bar graphs are showing level of **D:** plasma cholesterol level [mg/dL], **E:** blood glucose levels [mg/dL] and **F:** Triglycerides [mg/dL]. Data are presented as mean ± SEM and two-way ANOVA was performed with Sidak’s multiple comparison post-hoc test (*p < 0.05). N= 6 for each group.

As anticipated, rAAV8-PCSK9^D377Y^ administration resulted in a strong reduction in hepatic LDL receptor protein levels compared with saline-injected mice after both 12 and 20 weeks of intervention (**Figure 1B, C**). All mice thrived well, and body weight (**Supplementary Table 1**), blood lipids and glucose levels differed among treatment groups (**Figure 1D-E**). Plasma cholesterol levels were higher in the HGHCi HFD group than in the HCi HFD animals both at the early and late time points (**Figure 1D**); however, plasma triglyceride levels were higher in the HGHCi HFD vs. HCi HFD group only after 20 weeks (**Figure 1F**). Plasma triglyceride, cholesterol and glucose levels were comparable between rAAV8-PCSK9^D377Y^ (HCi HFD group) and LDLR KO mice on a HFD (**Figure 1D-F**).

Analyses of hematoxylin and eosin (H&E)- and Oil Red O-stained aortic root sections as well as the assessment of the aortic plaque score (**Supplementary Figure III A**) showed larger atherosclerotic plaques at both vascular sites (aortic sinus and BCA) in HGHCi HFD mice than in HCi HFD mice, both at early and late time points (aortic sinus at 12 weeks: HCi HFD 16% vs. HGHCi HFD 37% plaque area; 20 weeks: HCi HFD 29% vs. HGHCi HFD 43% plaque area) (**Figure 2A-D**). Control HFD mice had no lesions, whereas all rAAV8-PCSK9^D377Y^-injected mice on HFD (HCi HFD group) developed atherosclerosis to a similar extent as LDLR KO mice on HFD after 12 weeks and with no significant difference after 20 weeks (**Figure 2A-I**). In addition, HGHCi mice on a HFD had larger Oil Red O-stained plaques in the aortic sinus than LDLR KO HFD mice at both time points (**Figure 2C, D**). There was no significant difference in atherosclerotic plaque size between HGHCi HFD and LDLR KO HFD (**Figure 2E, F**) in the truncus brachiocephalic artery (BCA). In contrast to the aortic sinus, we found no significantly increased lesion area in the truncus brachiocephalic artery at 12 weeks, suggesting that the lesion develops earlier in the aortic sinus than in the brachiocephalic artery (**Figure 2E, F**). Image analysis of cross sections of BCA revealed a comparable EEL and IEL-area in HCi HFD and LDLR HFD mice, but thicker media and reduced lumen area in LDLR HFD mice (**Figure 2G-J**), the latter in line with increased plaque size in BCA of the LDLR HFD group (**Figure 2E, F**). In HGHCi HFD mice, EEL, IEL, media, and lumen area were reduced at 12 weeks compared with the HCi-HFD group. After 20 weeks, a reduction in lumen area persisted in HGHCi HFD mice compared with HCi HFD mice (**Figure 2J**), whereas EEL, IEL and media were comparable between the two groups (**Figure 2G-I**).

**Figure 2:**
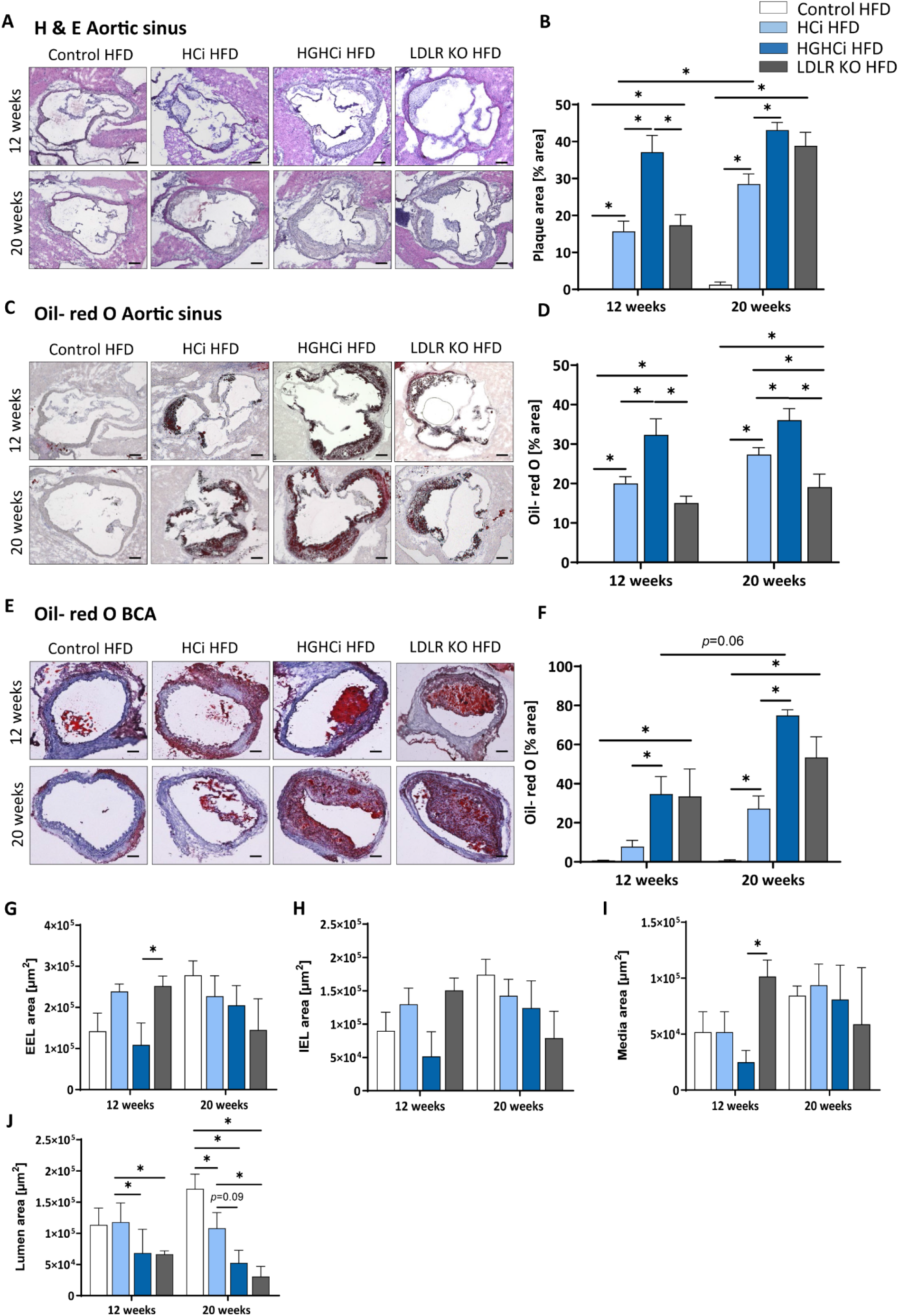
Mice with combined hyperlipidemia and hyperglycemia show larger plaques than hyperlipidemic mice. **A-D**: Representative histological images showing cross-sections of aortic sinus stained with Hematoxylin/ Eosin (H & E, **A**) and bar graphs summarizing data (**B**). Representative histological images showing aortic sinus sections stained with Oil-red O (**C**) and bar graphs summarizing data (**D**). **E-H:** Representative histological images showing truncus brachiocephalic arteries (BCA) sections stained with Oil-red O (**E**) and bar graphs summarizing data (**F**). Stainings were imaged at 4 × magnification (scale bar 100 μm) (**A, C, E**). HFD control 12 and 20 weeks (N= 7), HCi HFD 12 and 20 weeks (N= 5), and HGHCi HFD 12 and 20 weeks (N= 6). Determination of the external elastic lamina (EEL) (**G**), internal elastic lamina (IEL) (**H**), media area (**I**) and the lumen area (**J**) from Oil-red O stained cross-sections of the BCA (see Supplementary Figure II). Data presented as mean ± SEM and two-way ANOVA were performed with Sidak’s multiple comparison post-hoc test (*p < 0.05). Control HFD: Wild type mice without rAAV8-PCSK9^D377Y^ injection on high fat diet (HFD); HCi HFD: rAAV8-PCSK9^D377Y^ injection plus HFD (hyperlipidemic); HGHCi HFD: rAAV8-PCSK9^D377Y^ and streptozotocin injection and HFD (hyperlipidemic and hyperglycemic)

### Less stable plaque phenotypes in hyperglycemia and hyperlipidemia (HGHCi) versus hyperlipidemic (HCi) mice

In addition to plaque size, plaque stability is an important determinant of clinical outcome. We therefore evaluated indirect parameters of plaque stability in HCi, HGHCi and LDLR KO mice. Indeed, signs of plaque instability were more pronounced in HGHCi HFD mice than in HCi HFD mice, as evidenced from an increased necrotic core area, thinner fibrous caps, and reduced intraplaque α-SMA-positive cells, while no changes in the total vessel lumen were observed (**Figure 3A-J, Supplementary Figure IIIB**). Measurement of indirect markers of plaque stability was analyzed in Movat pentachrome-stained cross-sections of the aortic sinus (**Figure 3A-C**) and truncus brachiocephalic artery (**Figure 3D-F**). While HCi HFD and LDLR KO HFD mice developed few atherosclerotic lesions with increased necrotic core areas in the aortic sinus at 12 weeks (Figure 3A), diabetic HGHCi HFD mice showed increased necrotic core areas at this early time point. At 20 weeks, diabetic HGHCi HFD mice had significantly increased necrotic core areas and thinner fibrous caps at both vascular sites (aortic sinus and truncus brachiocephalic artery) compared to HCi HFD and LDLR KO HFD mice (**Figure 3A-F**). Plaque morphology and stability depend in part on the cellular composition of plaques. After 20 weeks, the number of α-SMA-positive cells within plaques of the aortic sinus and truncus brachiocephalic artery of HGHCi HFD mice was significantly decreased compared to that of nondiabetic HCi HFD mice but not of LDLR KO HFD mice (**Figure 3H, J**). There was no difference in α-SMA in the aortic sinus between HGHCi HFD and HCi HFD and LDLR KO mice after 12 weeks (**Figure 3G, H**). In the truncus brachiocephalic artery, HCi HFD mice showed no increased plaque development at 12 weeks, and accordingly, the number of α-SMA-positive cells was very low in this group (**Figure 3I, J**). However, in all other groups (HGHCi and LDLR KO at 12 weeks and HCi, HGHCi, and LDLR KO at 20 weeks), the number of α-SMA-positive cells within plaques was significantly increased compared to the control group in BCA (**Figure 3I, J**). Furthermore, we investigated intraplaque hemorrhage (by the erythrocyte marker protein Ter-119) in HGHCi HFD mice and found no significant differences between the HCi HFD and HGHCi HFD groups (**Supplementary Figure IV**).

**Figure 3:**
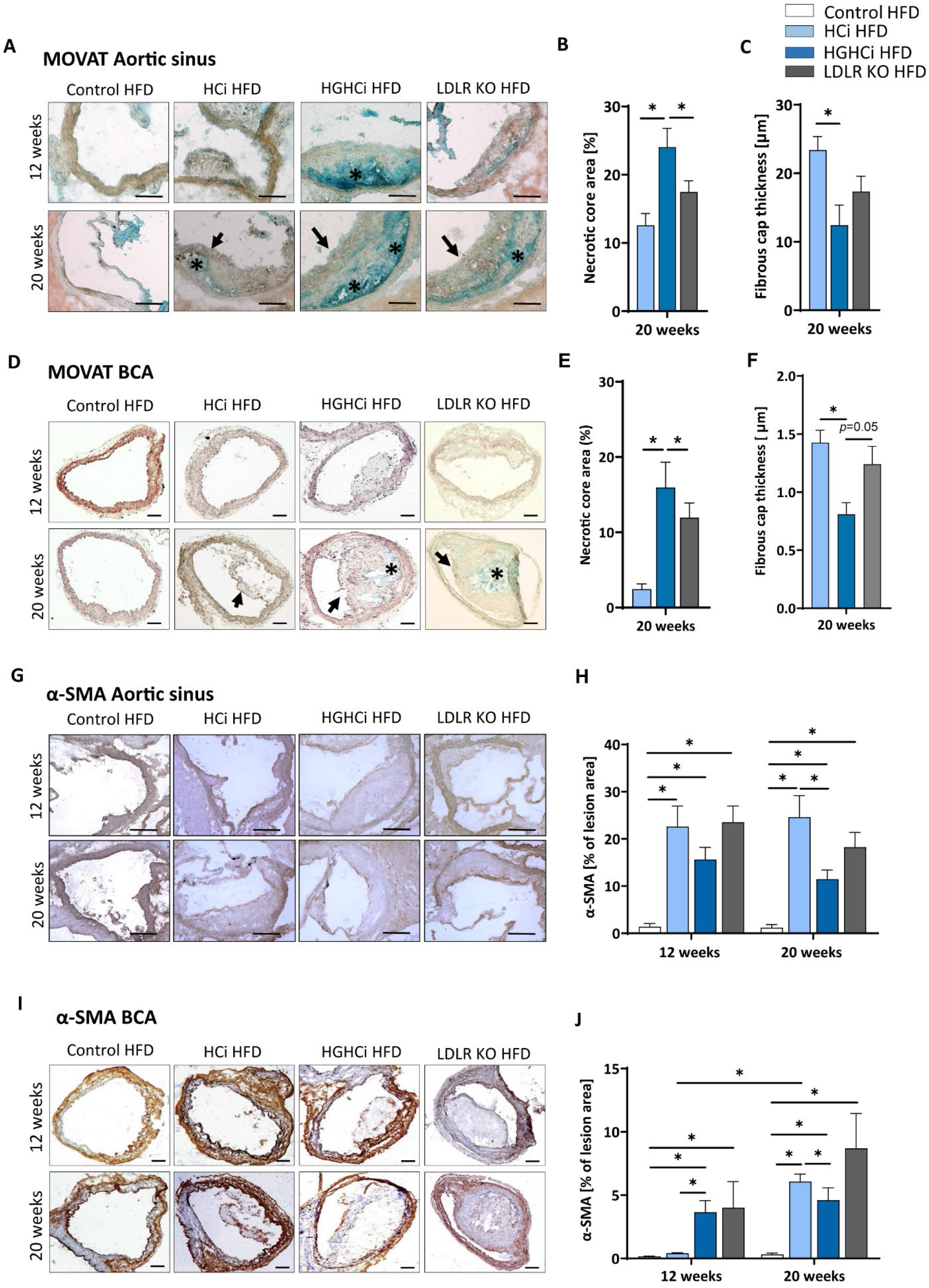
Increased necrotic core, reduced fibrous cap thickness, and α-SMA positive cells within plaques of hyperlipidemic and hyperglycemic mice as compared to hyperlipidemic mice. **A-C:** Representative images showing MOVATs pentachrome staining of cross-sections of the aortic sinus (**A**, scale bar 100 μm, 10x magnification). Bar graphs summarizing morphometric analyses of MOVATs stained images for necrotic core area (indicated by *) (**B**) and fibrous cap thickness (indicated by a black arrow) (**C**). Necrotic core area is depicted as % of total lesion area. **D-F:** Representative images showing MOVATs pentachrome staining of cross-sections of truncus brachiocephalic arteries (BCA) (**D**, scale bar 200 μm, 4x magnification). Bar graphs summarizing morphometric analyses of MOVATs stained images for necrotic core area (indicated by *) (**E**) and fibrous cap thickness thickness (indicated by a black arrow) (**F**). Data presented as mean + SEM and one-way ANOVA was performed with Sidak’s multiple comparison post-hoc test (*p < 0.05). **G,H:** Representative images showing immunohistochemical staining for α-SMA positive cells detected by HRP-DAB reaction (brown) on cross-sections of aortic sinus (**G**, **H**) (scale bar 100 μm, 10x magnification) and truncus brachiocephalic arteries (**I**, **J**) (scale bar 200 μm, 4x magnification). Corresponding bar graphs summarizing data (**H, J)**. Data presented as mean + SEM and two-way ANOVA were performed with Sidak’s multiple comparison post-hoc test (*p < 0.05). HFD control 12 and 20 weeks (N= 7), HCi HFD 12 and 20 weeks (N= 5), and HGHCi HFD 12 and 20 weeks (N= 6). Control HFD: Wild type mice without rAAV8-PCSK9^D377Y^ injection on high fat diet (HFD); HCi HFD: rAAV8-PCSK9^D377Y^ injection plus HFD (hyperlipidemic); HGHCi HFD: rAAV8-PCSK9^D377Y^ and streptozotocin injection and HFD (hyperlipidemic and hyperglycemic).

### Increased IL-1β plasma levels and expression of proinflammatory markers in hyperlipidemic and hyperglycemic mice compared to hyperlipidemic mice

As atherosclerosis is a chronic inflammatory disease and cytokines strongly influence disease development, we measured the plasma level of IL-1β and found significantly increased IL-1β levels in the diabetic HGHCi HFD group (5.8 pg/mL at 12 weeks; 5.6 pg/mL at 20 weeks) compared to the HCi HFD (1.9 pg/mL at 12 weeks; 1.8 pg/mL at 20 weeks) and LDLR KO HFD mice (3.5 pg/mL at 12 weeks; 2.6 pg/mL at 20 weeks) at both study time points (**Figure 4A**), indicating that systemic inflammation is already increased during early plaque development. As IL-1β was already increased after 12 weeks in the HGHCi HFD group, we analyzed the expression of marker genes of macrophage polarization (*Cd68* and *iNos* as M1 macrophage polarization marker genes; *Arg1* and *Fizz* as M2 marker genes) in the aorta of 12-week-old mice and found increased expression of M1 macrophage markers (*Cd68* and *iNos*) (**Figure 4B, C**) and correspondingly lower gene expression of M2 macrophage markers (*Arg1* and *Fizz*) (**Figure 4D, E**). In line with these results, the content of MOMA-2-positive cells was increased in plaques of the truncus brachiocephalic artery (BCA) (**Figure 4F, G**) of HGHCi HFD mice. In the aortic sinus, the number of MOMA-2-positive cells was elevated in all groups exhibiting atherosclerosis (HCi, HGHCi, and LDLR KO at 12 and 20 weeks) (**Figure 4H, I**).

**Figure 4:**
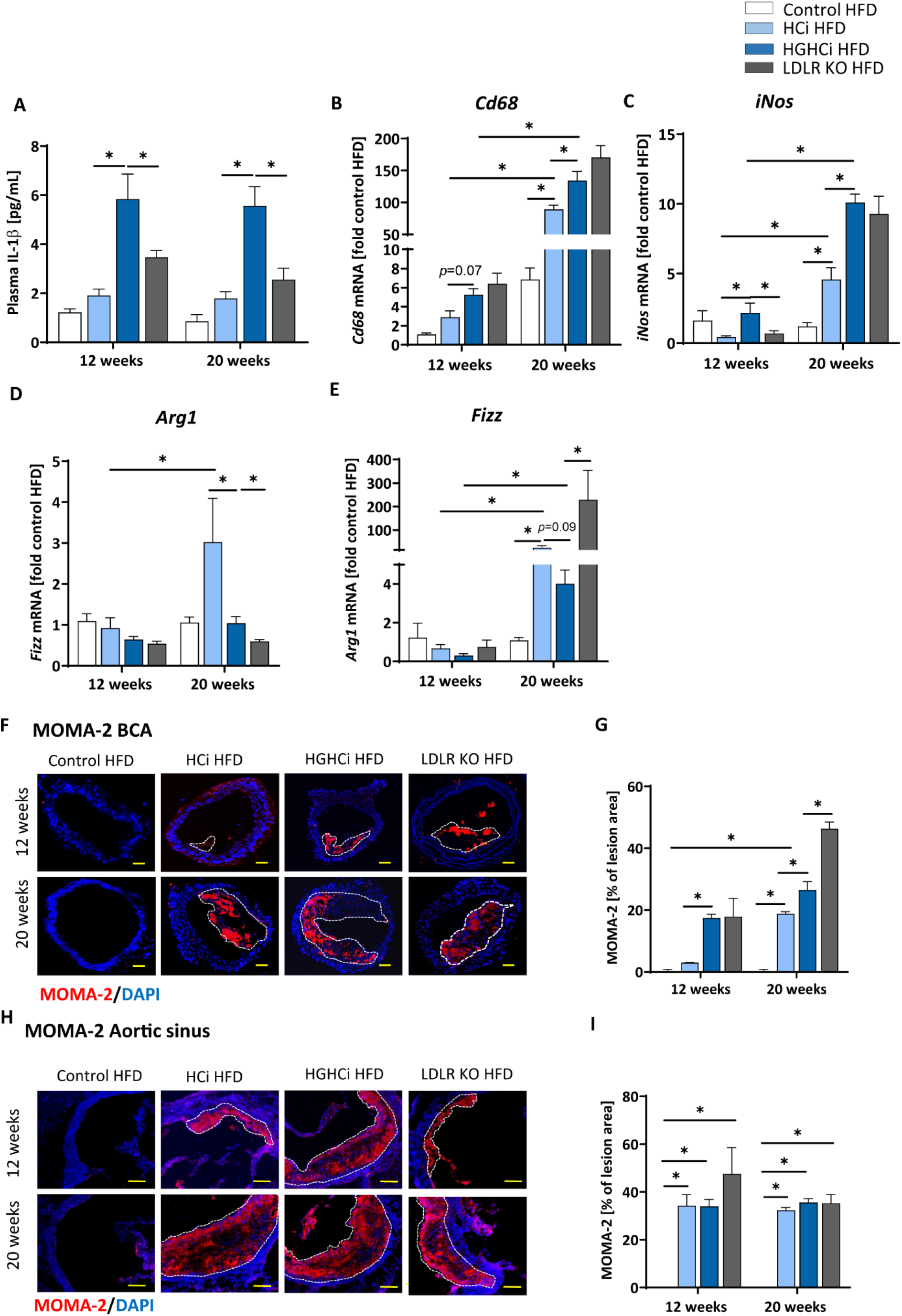
Increased IL-1β plasma level and expression of pro-inflammatory markers in hyperlipidemic and hyperglycemic mice as compared to hyperlipidemic mice. **A:** Plasma IL-1β [pg/mL] level at 12 and 20 weeks. **B-E**: mRNA expression of macrophage polarization markers. M1 macrophage polarization is depicted by *CD68* (**B**) and *iNos* (**C**) mRNA expression. *Arg1* (**D**) and *Fizz* (**E**) mRNA expression levels are shown as M2 polarization marker. The data are presented as mean ± SEM and were normalized on the mean of two housekeeping genes (*Hprt* and *B2m*). HFD control served as reference and was set at 1. **F, G:** Representative images showing immunofluorescence staining of truncus brachiocephalic arteries (BCA) sections for macrophage marker MOMA-2 (**F,** MOMA-2= red; DAPI nuclear counterstain= blue; plaque region is circled with a white dashed line, scale bar 200 μm, 4 × magnification) and bar graphs summarizing data (**G**). Representative images showing immunofluorescence staining of cross-sections of aortic sinus for MOMA-2 (**H,** MOMA-2= red; DAPI nuclear counterstain= blue; plaque region is circled with a white dashed line, scale bar 100 μm, 10 × magnification) and bar graphs summarizing data (**I**). Data presented as mean ± SEM and two-way ANOVA were performed with Sidak’s multiple comparison post-hoc test (*p < 0.05). HFD control 12 and 20 weeks (N= 7), HCi HFD 12 and 20 weeks (N= 5), and HGHCi HFD 12 and 20 weeks (N= 6). Control HFD: Wild type mice without rAAV8-PCSK9^D377Y^ injection on high fat diet (HFD); HCi HFD: rAAV8-PCSK9^D377Y^ injection plus HFD (hyperlipidemic); HGHCi HFD: rAAV8-PCSK9^D377Y^ and streptozotocin injection and HFD (hyperlipidemic and hyperglycemic).

Taken together, the observed shift toward increased lesion area, inflammation and necrotic core area and reduced fibrous cap thickness and α-SMA content suggests that plaque stability is reduced in HGHCi HFD mice compared to HCi HFD mice(20, 21).

### Identification of HGHCi-specific transcriptional responses in the aorta

We next aimed to identify transcriptional signatures that are engaged in the vasculature by combining chronic hyperlipidemia and hyperglycemia to generate hypotheses about signaling pathways contributing to larger but unstable plaques in HGHCi HFD mice. To this end, we conducted unbiased gene expression analyses (RNAseq) in the cohorts of control HFD, HCi HFD and HGHCi HFD mice at 12 weeks. In aortic tissue of HCi HFD mice, the expression of 942 genes was induced compared to the control mice solely on a HFD (**Figure 5A**). Conversely, gene expression in the aorta of HGHCi HFD mice was strikingly different from that of HCi HFD mice. Gene expression of a large set of genes was dysregulated in HGHCi HFD mice. Thus, in aortic tissue of HGHCi HFD mice, the expression of 2759 genes was induced compared to aortic samples of saline-treated HFD-fed mice (control HFD) (**Figure 5A**). A total of 418 genes showed similar expression in both HGHCi HFD and HCi HFD mice, and differential regulation of 2341 genes could be specifically assigned to combined hyperglycemia and hyperlipidemia.

**Figure 5:**
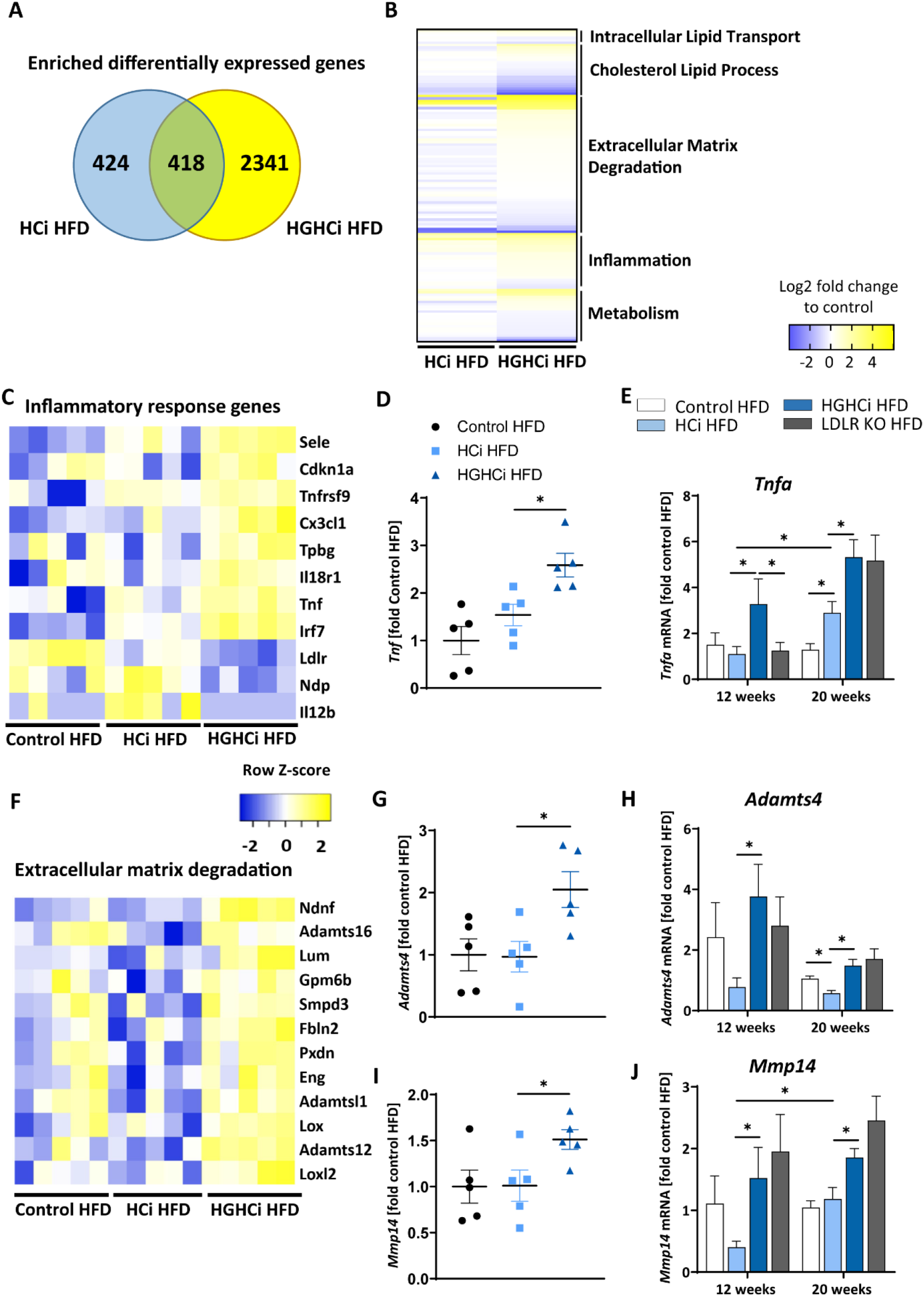
Combined hyperglycemia and hyperlipidemia induce large set of differentially expressed genes and activates inflammatory response and extracellular matrix degradation pathways. RNA sequencing in aortic tissue of control HFD, HCi HFD and HGHCi HFD mice after 12 weeks. Wild type mice without AAV8-PCSK9^D377Y^ injection on high fat diet (control HFD) or injected with rAAV8-PCSK9^D377Y^ and on HFD (hyperlipidemic, HCi HFD) or injected with both rAAV8-PCSK9^D377Y^ and streptozotocin and fed HFD (hyperlipidemic and hyperglycemic, HGHCi HFD). **A**: Venn diagram showing overlap of genes significantly changed in HGHCi HFD or HCi HFD mice in relation to gene expression in control HFD mice. **B:** Heat map summarizing differentially expressed gene (DEGs) identified by RNA sequencing and DEGs related biological processes using Gene Ontology and EnrichR analysis. **C**: Heat map showing list of DEGs related to inflammatory response (GO:0006954) using Gene Ontology and EnrichR analysis. **D:** *Tnf* expression level (FPKM normalized on Control HFD) from RNASeq at 12 weeks (N=5/ group) and its (**E)** qPCR validation in all groups at both time points (12 and 20 weeks). **F**: Heat map showing list of DEGs related to extracellular matrix degradation pathways (GO:0030198) using Gene Ontology and EnrichR analysis. RNA expression levels of selected ECM degradation genes *Adamts4* (**G, H**) and *Mmp14* (**I, J**) (FPKM normalized on Control HFD) from RNASeq at 12 weeks (N=5/ group) (**G, I**) and its qPCR validation (**H, J**) in all groups at both time points (12 and 20 weeks). Gene count values larger than the average control are represented in yellow, while lower counts than the average control are represented in blue. Whenever transcript values are close to the control value, samples are colored in white. The data are presented as mean ± SEM and qPCR data were normalized on the mean of two housekeeping genes (*Hprt* and *B2m*). HFD control served as reference control. Two-way ANOVA was performed with Sidak’s multiple comparison post-hoc test (*p < 0.05).

We next performed functional annotation analysis to study the gene pathways that are specifically engaged in the vessels of the HGHCi HFD group and observed that combined hyperglycemia plus hyperlipidemia led to significant changes in the expression of genes encoding signaling molecules involved in inflammation, intracellular lipid transport, cholesterol metabolic processes, extracellular matrix (ECM) degradation, and cellular metabolism (**Figure 5B**). We further performed gene ontology analysis on genes that were induced in aortic tissue of HCi HFD and HGHCi HFD mice. Gene ontology analysis of differentially expressed genes revealed that combined hyperglycemia and hyperlipidemia led to the upregulation of genes involved in the inflammatory response (GO:0006954) (**Figure 5C**) and ECM degradation (GO:0030198) (**Figure 5F**). We further investigated the expression of TNFa, which is a central mediator of inflammatory reactions and plays an important role in atherogenesis, and found that it was significantly upregulated in the HGHCi HFD mice compared to HCi HFD mice at both study time points (**Figure 5D, E**). Moreover, we analyzed additional matrix metalloproteinases, such as Mmp14 and Adamts4 (A Disintegrin and Metalloproteinase with Thrombospondin motifs 4), in the RNA-seq data and corroborated their dysregulation by qPCR analysis (**Figure 5G-J**). Both marker genes were upregulated in HGHCi HFD mice compared to HCi HFD mice after both 12 and 20 weeks (**Figure 5G-J**).

### The Paigen diet accentuates atherosclerosis and promotes plaque instability in mice with combined hyperglycemia and hyperlipidemia

We next tested the atherosclerosis model following induction of combined hyperglycemia and hyperlipidemia (HGHCi) in mice fed a Paigen diet (PD), which leads to higher plasma cholesterol levels (7) and might thus further exacerbate the formation of atherosclerotic plaques. PD-fed mice were treated with the same batch and dose of rAAV8-PCSK9^D377Y^, and LDLR KO mice on the PD served as a reference for comparison. Saline-injected mice fed the PD served as a control for rAAV8-PCSK9^D377Y^ intervention. Hyperlipidemic and hyperlipidemic plus hyperglycemic mice on the PD are hereafter called HCi PD and HGHCi PD mice, respectively, for simplicity. In HGHCi PD mice, we observed an increased mortality (25% in week 8, i.e., 5 out of 20 mice died in this group (**Supplementary Figure I, A**). Based on these incidences, it was decided that the remaining animals in this group should be sacrificed and analyzed 8-9 weeks after induction of combined hypercholesterolemia and hyperglycemia, since the 12- and 20-week time points chosen for the HGHCi group on HFD might not have been reached. Thus, mice from all groups on the PD diet were euthanized 8-9 weeks after rAAV8-PCSK9^D377Y^ injection to quantify atherosclerosis (**Figure 6A**). Hepatic LDL receptor protein expression was already abrogated at this time point compared with saline-injected mice (control PD) (**Figure 6B, C**). As expected, body weight (**Supplementary Table 1)**, plasma cholesterol levels and blood glucose levels differed among the HCi and HGHCi PD groups (**Figure 6D-F),** while HCi PD mice and LDLR KO PD mice did not differ regarding cholesterol, triglyceride and glucose levels (**Figure 6D-F**). In a preliminary pilot study, we compared the plasma cholesterol levels of the HGHCi PD group with STZ-treated mice (STZ PD) and citrate-treated controls (citrate PD) fed a Paigen diet (**Supplementary Figure V**). Notably, mice with STZ-evoked hyperglycemia and on a high-cholesterol, high-fat Paigen diet (without AAV-mPCSK9 treatment) did not exhibit an increase in plasma cholesterol (**Supplementary Figure IV).** Application of the Paigen diet exacerbated atherosclerosis in both HGHCi PD and HCi PD mice compared to the corresponding groups on the HFD diet (p< 0.05). Interestingly, the aortic root plaque sizes of HGHCi PD and HCi PD mice at 8 weeks (**Figure 6G-J)** were comparable to those of HGHCi and HCi mice on a HFD at 20 weeks (**Figure 2A**). Consistent with data from the HFD groups, signs of plaque instability were more pronounced in HGHCi PD mice than in HCi-PD mice as measured by decreased collagen deposition (picrosirius red staining) and increased plaque CD68-positive cells in HGHCi PD mice in comparison with HCi PD mice (**Figure 7A-D**). Taken together, these data suggest that the PD aggravates atherosclerosis in mice and shows plaque formation after 8 weeks, therefore representing a direct inducible diabetic atherosclerosis model with more aggravated atherosclerosis than HGHCi HFD.

**Figure 6:**
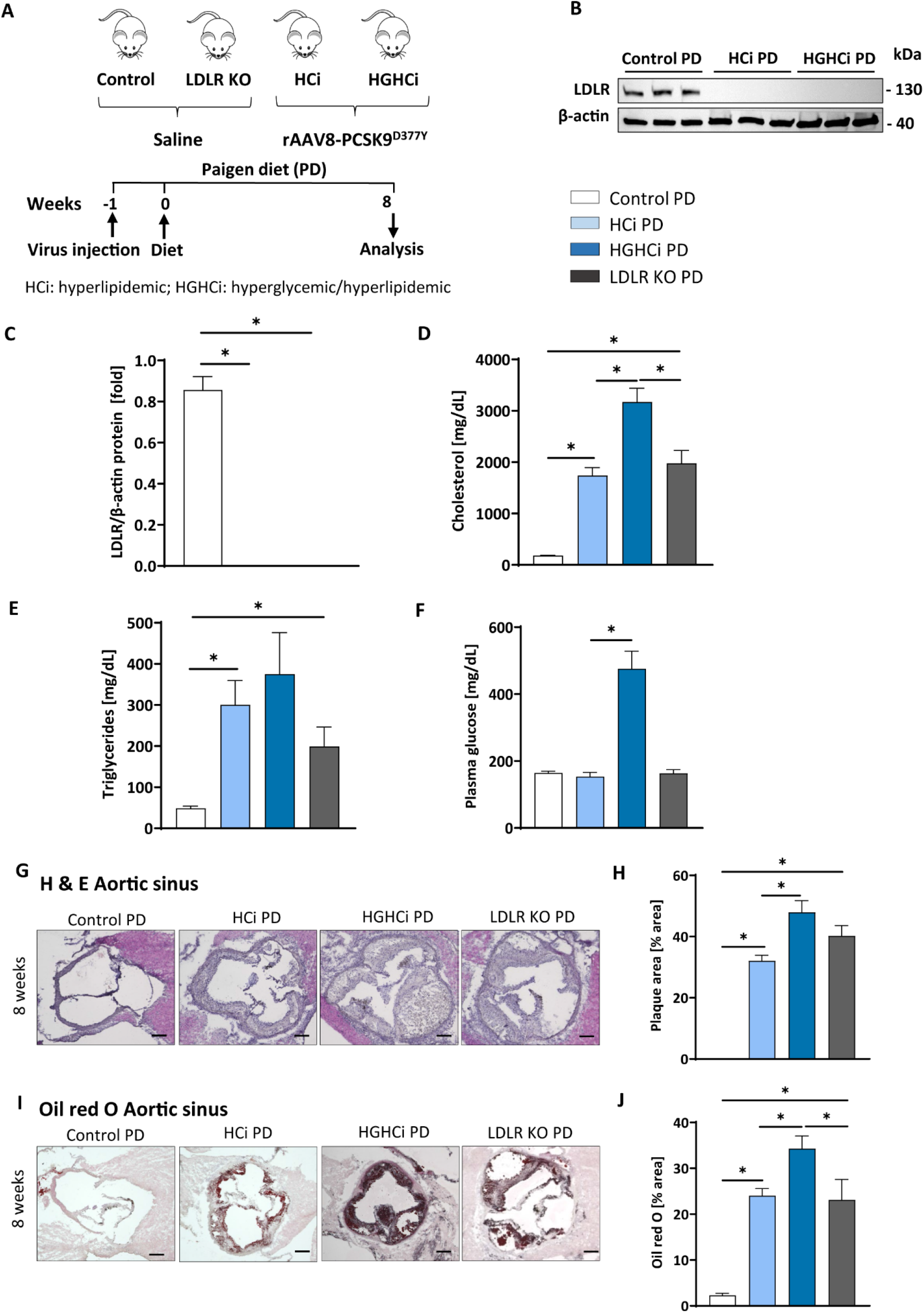
Paigen diet accelerates plaque formation after 8 weeks in the rAAV8-PCSK9 streptozotocin induced hyperglycemic atherosclerosis mouse model. **A**: Schematic summary of the experimental setup. Mice were analyzed after 8 weeks of interventions initiation. **B**: Representative immunoblot showing hepatic protein levels of low-density lipoprotein receptor (LDLR). β-actin was used as loading control and bar graphs summarizing data. Data were normalized on β-actin and control PD mice were set at 1 (**C**). Bar graphs summarizing data of plasma cholesterol level [mg/dL, **D**], triglycerides [mg/dL, **E**), and blood glucose levels [mg/dL, **F**]. **G-J:** Representative histological images showing aortic sinus sections stained with Hematoxylin Eosin (H & E, **G**) and bar graphs summarizing data (**H**). Representative histological images showing aortic sinus sections stained with Oil-red-O (**I**) and bar graphs summarizing data (**J**). Data presented as mean ± SEM and one-way ANOVA was performed with Sidak’s multiple comparison post-hoc test (*p < 0.05). Scale bar 200 μm. Control paigen diet (PD) (N=6), HCi PD (N=6), HGHCi PD (N=6), LDLR KO PD (N=5). Control PD: Wild type mice without rAAV8-PCSK9^D377Y^ injection on PD; HCi PD: mice injected with rAAV8-PCSK9^D377Y^ and fed PD (hyperlipidemic); HGHCi PD: mice injected with both rAAV8-PCSK9^D377Y^ and streptozotocin and fed PD (hyperlipidemic and hyperglycemic); LDLR KO PD: LDLR KO mice fed PD.

**Figure 7:**
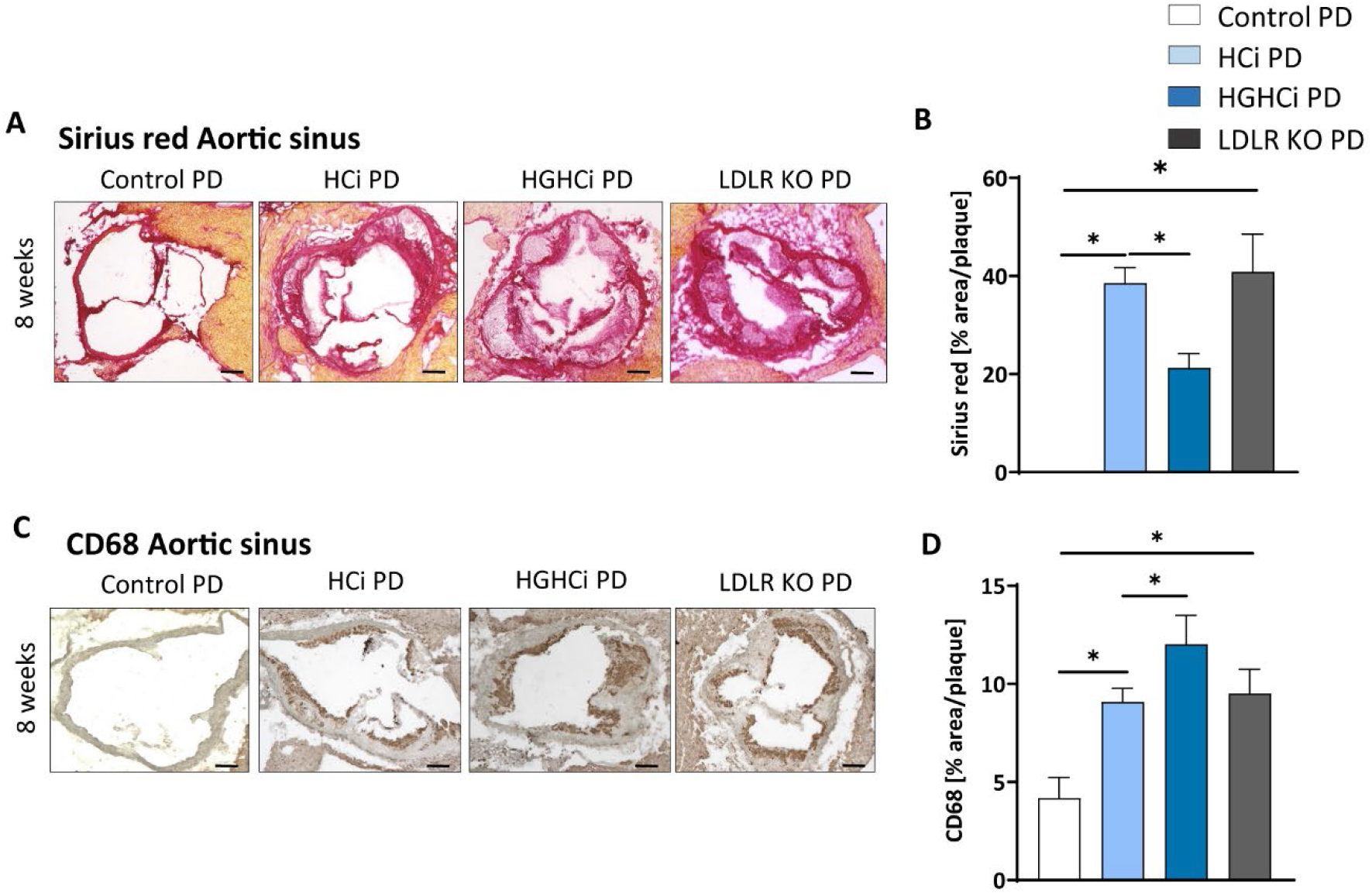
Hyperlipidemic and hyperglycemic mice fed a Paigen diet (PD) show less collagen and increased inflammation within aortic plaques compared to hyperlipidemic mice. **A**: Representative images showing picrosirius red staining for collagen in aortic sinus sections and bar graphs summarizing data (**B**). Scale bar 200 μm. **C:** Representative images showing immunohistochemical staining of aortic sinus sections for macrophages marker CD68 (positive cells detected by HRP-DAB reaction, brown) and bar graphs summarising data (**D**). Data presented as mean ± SEM and one-way ANOVA was performed with Sidak’s multiple comparison post-hoc test (*p < 0.05). Scale bar 100 μm. Control paigen diet (PD) (N= 7), HCi PD (N= 4), HGHCi PD (N=4), LDLR KO PD (N=5). Control PD: Wild type mice without rAAV8-PCSK9^D377Y^ injection on PD; HCi PD: mice injected with rAAV8-PCSK9^D377Y^ and fed PD (hyperlipidemic); HGHCi PD: mice injected with both rAAV8-PCSK9^D377Y^ and streptozotocin and fed PD (hyperlipidemic and hyperglycemic); LDLR KO PD: LDLR KO mice fed PD.

## Discussion

The most frequently used models to study diabetes mellitus-associated atherosclerosis in mice rely on genetically modified models. Thus, mouse studies evaluating diabetes-associated atherosclerosis rely on ApoE^-/-^ or LDLR^-/-^ mice, where hypercholesterolemia is combined with chronic hyperglycemia following beta-cell destruction by injection of streptozotocin (STZ) or viral infection (22) or by crossbreeding with mouse strains carrying a point mutation in the gene encoding insulin leading to a misfolding of the proinsulin 2 protein (Ins2^+^/Akita) (4, 23). Induction of hyperglycemia with STZ led to higher plasma cholesterol levels and showed significant acceleration of atherosclerotic lesion formation in ApoE^-/-^ and LDLR^-/-^ mice compared to nondiabetic ApoE^-/-^ and LDLR^-/-^ controls (6, 24–30). Additionally, the severe hyperglycemia in Ins2^+^/Akita mice crossbred with ApoE^-/-^ and LDLR^-/-^ leads to a pronounced increase in atherosclerosis and non-HDL cholesterol and triglyceride levels compared to those of nonhyperglycemic control mice (4, 31, 32). Although these genetic models are useful tools for studying diabetic complications, intensive crossbreeding of these mouse lines is required. This becomes even more laborious and time-consuming when diabetic long-term complications are to be induced in knockout mouse lines for evaluation of the causal role of the gene of interest under certain metabolic conditions. Concomitant induction of both hypercholesterolemia and hyperglycemia at discrete time points in a given mouse line has not been reported.

Here, we describe a novel strategy to induce an atherosclerosis model aggravated by concomitant induction of hyperglycemia without the necessity of genetic germline engineering but via rAAV8-mediated gene transfer of mutant PCSK9^D377Y^ in hepatocytes in combination with STZ-evoked beta-cell destruction (HGHCi). Comparison of plasma cholesterol levels shows increased cholesterol levels in the HCi model, which increase further by additional induction of hyperglycemia in the HGHCi model. The additional increase in cholesterol levels is a consequence of the two-hit model because induction of hyperglycemia alone does not increase plasma cholesterol levels. The increased plasma cholesterol levels in the diabetic mice (HGHCi group) are part of the phenotype of our model and resemble findings in previous studies (7,26,28,29). Dyslipidemia aggravated by hyperglycemia is independent of whether the diabetes was induced by STZ injection or genetically (e.g., InsAkita mutant mice). Increased cholesterol levels in the diabetic mice most likely reflect decreased lipoprotein clearance, and as such, they may reflect an important feature of diabetes-associated dyslipidemia. Thus, Goldberg et al. demonstrated that the plasma cholesterol levels of STZ-induced diabetic LDLR^-/-^ mice were twice those of nondiabetic control mice. The authors observed an increase in both VLDL and LDL. VLDL in plasma was more enriched in cholesterol, and both VLDL and LDL had high levels of ApoE (30). Further lipidomics analyses are required to define the lipid profile in diabetic and nondiabetic mice with PCSK9^D377Y^-induced dyslipidemia. We cannot differentiate whether the increased cholesterol level or other factors (e.g., the elevated systemic inflammation) or their combination is responsible for the increased plaque progression in HGHCi mice compared to HCi mice. However, STZ treatment alone did not lead to elevated LDL cholesterol in our study (Supplementary Figure V) or in previous studies (6), where it was shown that diabetic mice are resistant to atherosclerosis even in the presence of a high-fat diet. To address the question of whether the additional increase in LDL cholesterol (evoked by hyperglycemia in PCSK9^D377Y^-treated mice) is indeed responsible for increased plaque progression, one might design a study using a selective LDL-lowering therapy without an effect on hyperglycemia (e.g., by HMG-CoA reductase inhibitors) in the HGHCi model. The influence of systemic inflammation on increased plaque progression in the HGHCi group compared to the HCi group could be investigated with an anti-inflammatory therapy (e.g., by the anti-IL-1β antibody canakinumab). Studies addressing such mechanistic links will be required in the future.

Furthermore, we show that combined hyperlipidemia and hyperglycemia led to a significant enhancement of atherosclerotic plaque formation compared to mice in which hypercholesterolemia was induced without hyperglycemia (HCi). The HCi model in rAAV8-mediated mutant PCSK9^D377Y^-expressing mice leads to downregulation of hepatic LDL receptor expression and the development of atherosclerotic lesions. The lesion phenotypes closely resemble those in the established LDL-receptor knockout mouse model. Mice in the HGHCi treatment group displayed larger plaques than norrmoglycemic HCi mice on the same diet and LDLR knockout mice (after 12 weeks). Treatment-induced systemic atherosclerosis (e.g., in the aortic sinus and in the brachiocephalic artery) and inflammation were measured by increased plasma levels of IL-1β. The plaque phenotypes induced by HGHCi compared to HCi mice are characterized by increased necrotic core area and decreased fibrous cap thickness. Plaques are characterized by increased gene expression of M1 macrophage markers and MOMA-2-positive cells as well as a reduced number of α-SMA-positive cells (at 20 weeks). A comparably increased amount of α-SMA-positive cells was observed in HGHCi HFD and LDLR KO HFD mice at 12 weeks in the aortic sinus and BCA, which is characteristic of early lesion development (33, 34). In HCi HFD mice, the number of α-SMA-positive cells was also increased in the aortic sinus after 12 weeks but not in the BCA, where a small plaque size was observed at that time point. The reason for these differences in plaque size and α-SMA-positive cell content between the aortic sinus and BCA in HCi mice after 12 weeks can be explained by an earlier development of lesions at the aortic sinus, which has been described in other studies (35–37). Whether the earlier onset of α-SMA-positive cell proliferation and migration in the BCA of HGHCi HFD and LDLR KO mice is due to stronger systemic inflammation indicated by the higher increase in IL-1β levels compared with HCi HFD mice at 12 weeks needs to be investigated in future studies. Taken together, these data show that the novel HGHCi model provides an overview of plaque development and progression at two locations in the aorta to study interventions aiming to improve plaque size and, importantly, plaque stability. A similar severity of atherosclerosis was observed with the Paigen diet, albeit much earlier, at 8 weeks post-intervention initiation, compared to 20 weeks with the HFD. It has been reported that the addition of cholate in combination with a high cholesterol and high fat content in the Paigen diet boosts hypercholesterolemia by facilitating fat and cholesterol absorption, resulting in very early fatty streak lesions in the aortic root and proximal aorta in C57BL/6 mice (38). Thus, our findings are in line with previous work in genetic models, demonstrating that hyperglycemia promotes unstable plaques in both ApoE^-/-^ and LDLR^-/-^ mice (14, 39–41). Our results identify the HGHCi PD model using a Paigen diet as a novel option for specific experimental setups where strong plaque formation and progression are needed at an early time point.

Our unbiased gene expression analysis revealed that combined hyperlipidemia and hyperglycemia (HGHCi) dysregulates a large set of genes (i.e., 2341 genes) in the atherosclerotic aorta compared to hyperlipidemia alone (HCi). The aortic tissue gene expression profile of HGHCi mice fed a HFD differed markedly from HCi HFD mice. Among the pathways that were most prominently affected were the inflammatory response, ECM degradation and metabolism. Upregulated proinflammatory genes, such as *Cx3cl1* (or fractalkine) and *Il18r1*, have been shown to be involved in atherogenesis. *Cx3cl1* is a chemokine and exerts cytotoxic effects on the endothelium. Its membrane-bound form promotes adhesion of rolling leukocytes onto the vessel wall, while in its soluble form, it serves as a potent chemoattractant for CX3CR1-expressing cells. Therefore, it affects the context and stability of the atherosclerotic plaque. Blocking the CX3CL1/CX3CR1 pathway in *in vivo* studies ameliorated the severity of atherosclerosis (42). Il18r1 is a member of the interleukin receptor family and is expressed on T-lymphocytes but also on cell types associated with atherogenesis, such as macrophages, endothelial cells, and smooth muscle cells (43). After binding its ligand IL-18, it leads to the expression of IFNγ via the NF-kB-mediated signaling pathway (44). High expression of IL18R is found in plaque-resident macrophages and endothelial cells in humans (45).

The increased expression of genes related to ECM degradation (e.g., *Mmp14* and *Adamts4*) is consistent with previous findings showing that glucose-induced advanced glycation end products (RAGE) mediate modification of the components of the ECM and accelerate atherosclerosis under diabetic conditions (46–49). The ECM degradation genes *Mmp14* and *Adamts4* were shown to play key roles in plaque stability. Loss of *Adamts4* in ApoE KO mice increased plaque stability (50), and ADAMTS4 expression in humans was upregulated during carotid atherosclerotic plaque development. Furthermore, ADAMTS4 serum levels were associated with increased plaque vulnerability (51). Metalloproteinases (MMPs), such as *Mmp14*, play an important role in the pathogenesis of atherosclerosis by participating in vascular remodeling, smooth muscle cell migration, and plaque disruption. Mmp14 is recognized as a prominent member of this family, causes pericellular degradation and has the ability to activate other matrix metalloproteinases. It is further detected at high levels in atherosclerotic plaques (52).

Genes involved in inflammation, such as *Tnfa, Cd68* and *iNos*, were shown to be upregulated in HGHCi HFD mice compared to HCi HFD mice, which has been described in previous studies where their expression is one of the key drivers in atherosclerosis-associated chronic inflammation in arterial blood vessels (48). The progression of atherosclerosis correlates directly with a local increase in TNF-α production in atherosclerotic plaques and blood (53). The expression data may provide a helpful resource for researchers using this model and suggest that this nongenetic model will be suitable for studying important aspects of the emerging crosstalk between inflammatory signaling, glucose metabolism and lipoprotein metabolism (23, 54–56).

### Limitations of the study

In this study, only male mice were analyzed. As reported previously, rAAV8-PCSK9^D377Y^-evoked hypercholesterolemia was 3-fold lower in females than in males (57), and consequently, the extent of atherosclerosis development can be expected to be lower or delayed. To characterize the development of atherosclerosis in our HGHCi model in females, new studies will be required in the future to elucidate sex-specific effects in this model. The aim of the study was to develop and characterize novel models of hyperlipidemia and hyperglycemic-associated atherosclerosis. Plasma lipid levels were not identical between groups. Due to the study design, a cohort with only STZ injection was not included, as hyperglycemia induced by STZ treatment or lymphocytic choriomeningitis virus (LCMV) infection alone is not sufficient to induce atherosclerosis development in mice (29, 39). Hence, conclusions about the impact of isolated hyperglycemia on the atherosclerotic plaque phenotype cannot be drawn from the current study.

Atherosclerosis development in mice and humans differs in several parameters, including lipoprotein metabolism. In mice, cholesterol is transported mainly in HDL particles due to a lack of cholesteryl ester transfer protein (CETP) expression, whereas humans show high CETP expression, which results in high LDL cholesterol levels (23). Furthermore, wild-type mice have significantly lower total cholesterol levels than humans, which may explain why they do not develop atherosclerosis under natural conditions. In recent years, atherosclerosis-associated genes have been identified and used to generate genetically modified mouse models that are used in atherosclerosis research. The two most commonly used genetic models in atherosclerosis research are the Apoe-/- and Ldlr-/- models (58, 59). Among other parameters, the models exhibit critical differences in cholesterol and lipid metabolism, which makes their use highly dependent on the scientific question. Apolipoprotein E (apoE) binds to chylomicrons and VLDL in plasma and acts as a ligand that initiates uptake of the remnants via the LDL receptor, thereby removing them from the circulation. Deletion of apoE in Apoe-/- mice results in an increase in cholesterol levels by the accumulation of chylomicrons and VLDL (58). In contrast, in the Ldlr-/- model, deletion of LDLR results in an increase in cholesterol levels by the accumulation of plasma LDL, which is much more comparable to the human situation than the elevation of plasma VLDL in Apoe-/- mice (23). Expression of the rAAV-PCSK9^D377Y^ (7) used in this study leads to increased degradation of LDLR, which is comparable to the deletion of LDLR in the Ldlr-/- model. Additionally, in this inducible model, there is an increase in total cholesterol due to the accumulation of LDL particles, and it was shown by Roche-Molina et al. (60) that the lipoprotein profiles of Ldlr-/- mice and rAAV-PCSK9^D377Y^-treated animals fed a HFD are very similar. Therefore, both the Ldlr-/- and the inducible rAAV-PCSK9^D377Y^ models are currently most suitable for cholesterol and lipid metabolism studies investigating processes in human atherosclerosis development (23). In addition to their lipoprotein profile, the models also differ from the human disease pattern with respect to the sites at which lesions develop. In humans, disease-relevant plaques develop preferentially in the coronary arteries, carotid arteries and peripheral arteries of the legs and arms, whereas in mouse models, lesions are mainly found in the aortic sinus, aortic arch and brachiocephalic artery (23). In addition, the extent of plaque development in mice is reduced, and large vulnerable plaques are often absent compared to humans; thus, plaque ruptures are rarely observed in genetic mouse models, which in turn occur in humans at advanced stages of atherosclerosis and lead to atherothrombotic vessel occlusions (23, 58, 59).

In the future, it may be possible to use the HGHCi model in preclinical animal models that more accurately reflect human lipoprotein metabolism and the development and progression of atherosclerotic lesions in coronary arteries than the already established genetic mouse models. Both STZ-mediated hyperglycemia in pigs and rabbits (61, 62) and PCSK9 gain-of-function overexpression-induced hypercholesterolemia in pigs (63) have been demonstrated to date and, in combination, may achieve further important disease-relevant findings about diabetes-accelerated atherosclerosis and inflammation in humans.

## Conclusion

We describe a novel nongenetically inducible mouse model using a defined treatment with gain-of-function PCSK9 viral particles, streptozotocin and high-fat diets. The detailed characterization of the plaque phenotypes and the expression analysis indicate that this new experimental approach can be used to study the clinically important pathogenesis of atherosclerosis in hyperglycemia and hyperlipidemia and can be induced at any given time point to prevent or therapeutically interfere with the development of atherosclerosis under diabetic conditions.

## Acknowledgement

We thank Sonja Talmon, Beate Hilbert, Christin Richter, Manuela Ritzal, Anja Barnikol-Oettler, and Kathrin Deneser for excellent technical support. We certify that all persons who have made substantial contributions to the manuscript, but who do not fulfill authorship criteria, are named in the acknowledgments section and have provided the corresponding author with written permission to be named in the manuscript.

## Authors contribution

**S.G.**, **K.S.** and **R.M**., interpreted the experimental work and prepared the manuscript. **A.W., R.M., S.G., I.G**. and **D.S.** performed and conducted the in vivo experiments; **S.G., I.G.,** and **C.M.,** performed histological analyses and figure preparation; **J.H., J.B**., and **H.K.** performed RNAseq analysis; **S.A.** and **S.F.** assisted in histology. **B.I.** and **U. L.** interpreted the data and assisted in manuscript preparation. **M.F.** designed the study and assisted in manuscript preparation.

## Data Availability

All data that support the findings of this study are available in this study within the manuscript and/or its supplementary materials.

## Supplemental Materials

**Supplementary Table 1.** Characteristics of mice cohort.

**Supplementary Figure I.** PD fed HGHCi mice showed higher mortality than HFD fed mice.

**Supplementary Figure II.** Determination of the external and internal elastic lamina, lumen area and media area.

**Supplementary Figure III.** Comparison of aortic plaque score and total vessel lumen area.

**Supplementary Figure IV.** Intraplaque hemorrhage is not increased in HGHCi HFD mice.

**Supplementary Figure V.** Hyperglycemia alone does not affect plasma cholesterol levels in mice fed a Paigen diet.

**Major resources table**

## Supplementary Material

**Supplementary Table 1.**
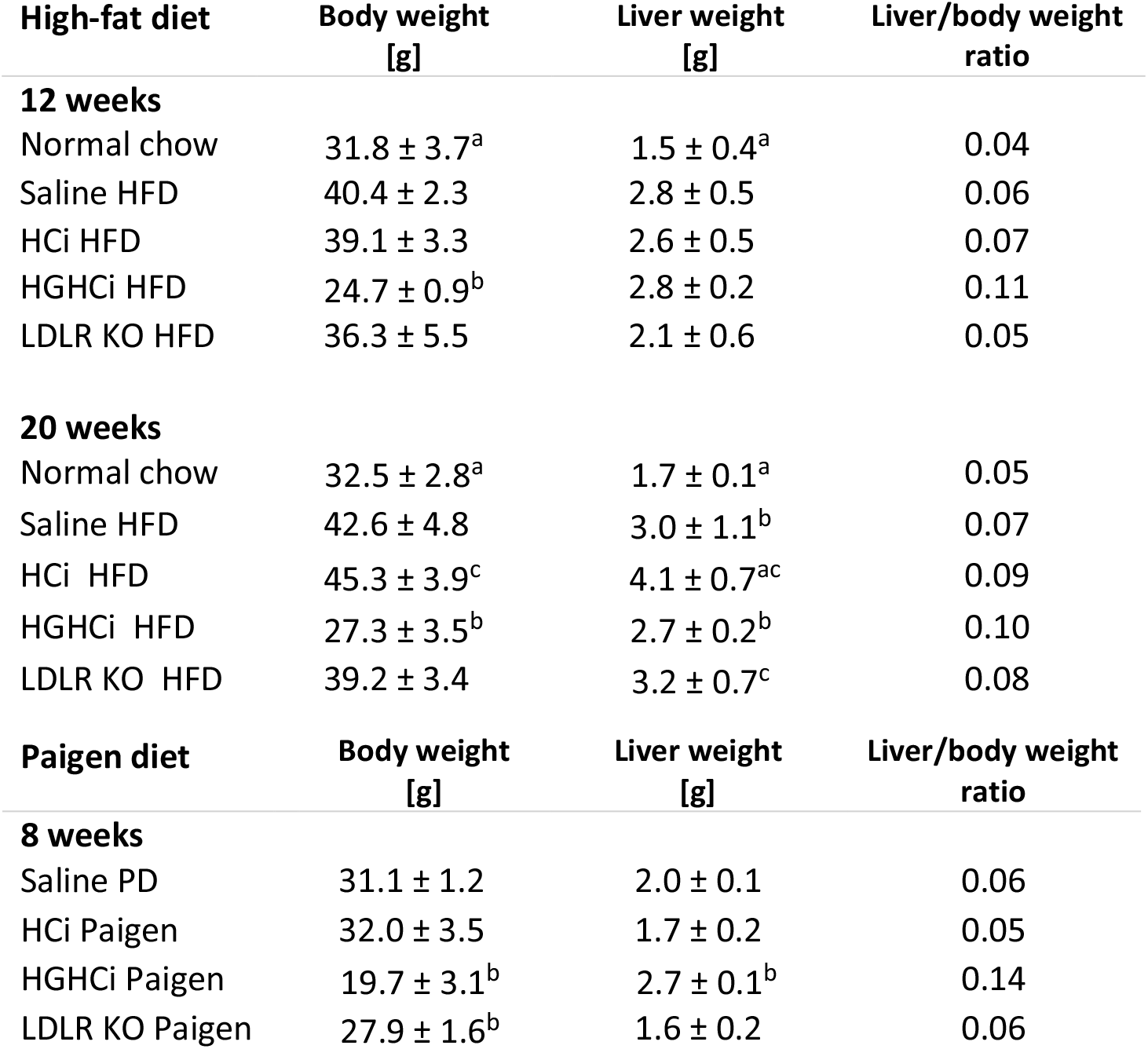
Characteristics of mice cohort. Data are shown as mean ± SEM and statistical comparisons between the groups were calculated using the one-way ANOVA and Sidak’s posthoc test. ^a^ p<0.05 vs Saline HFD/PD, ^b^ p<0.05 vs HCi HFD/PD, ^c^ p<0.05 vs 12 weeks. HFD: high-fat diet; HCi: hyperlipidemia model; HGHCi: hyperglycemia + hyperlipidemia model; PD: Paigen diet

**Supplementary Figure I.**
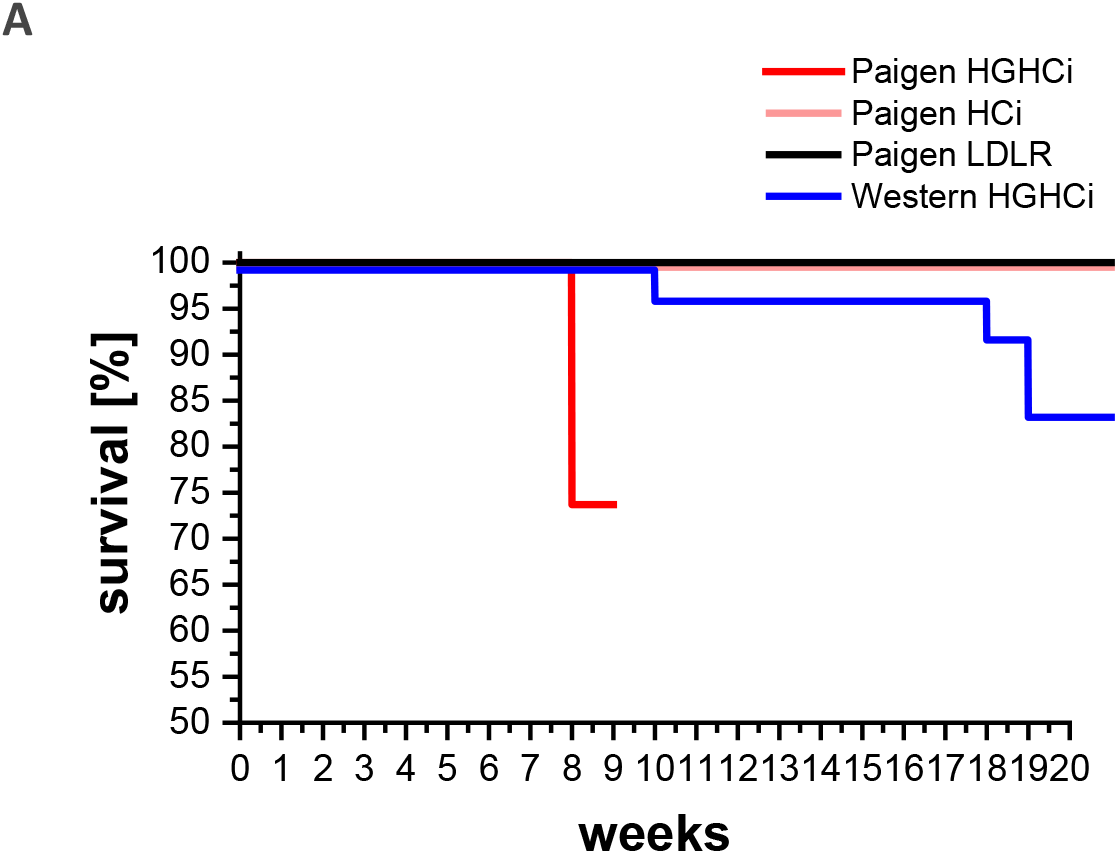
PD fed HGHCi mice showed higher mortality than HFD fed mice. **A:** Kaplan-Meier survival curves of HGHCi, HCi and LDL receptor KO mice on Paigen diet compared to HGHCi mice on Western diet (high-fat diet, HFD).

**Supplementary Figure II.**
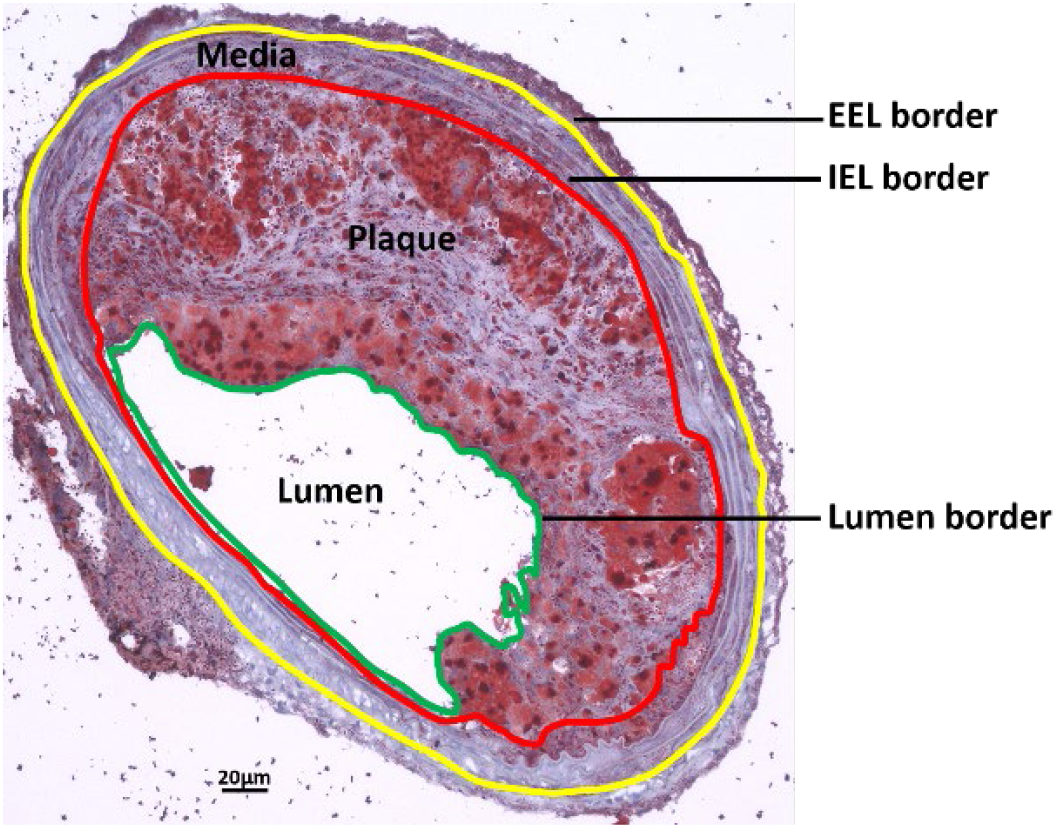
Determination of the external and internal elastic lamina, lumen area and media area. Representative image showing Oil-Red O stained BCA (**G**) for the measurement of the external elastic lamina (EEL, yellow line), internal elastic lamina (IEL, red line) and the lumen area (circled by a green line).

**Supplementary Figure III.**
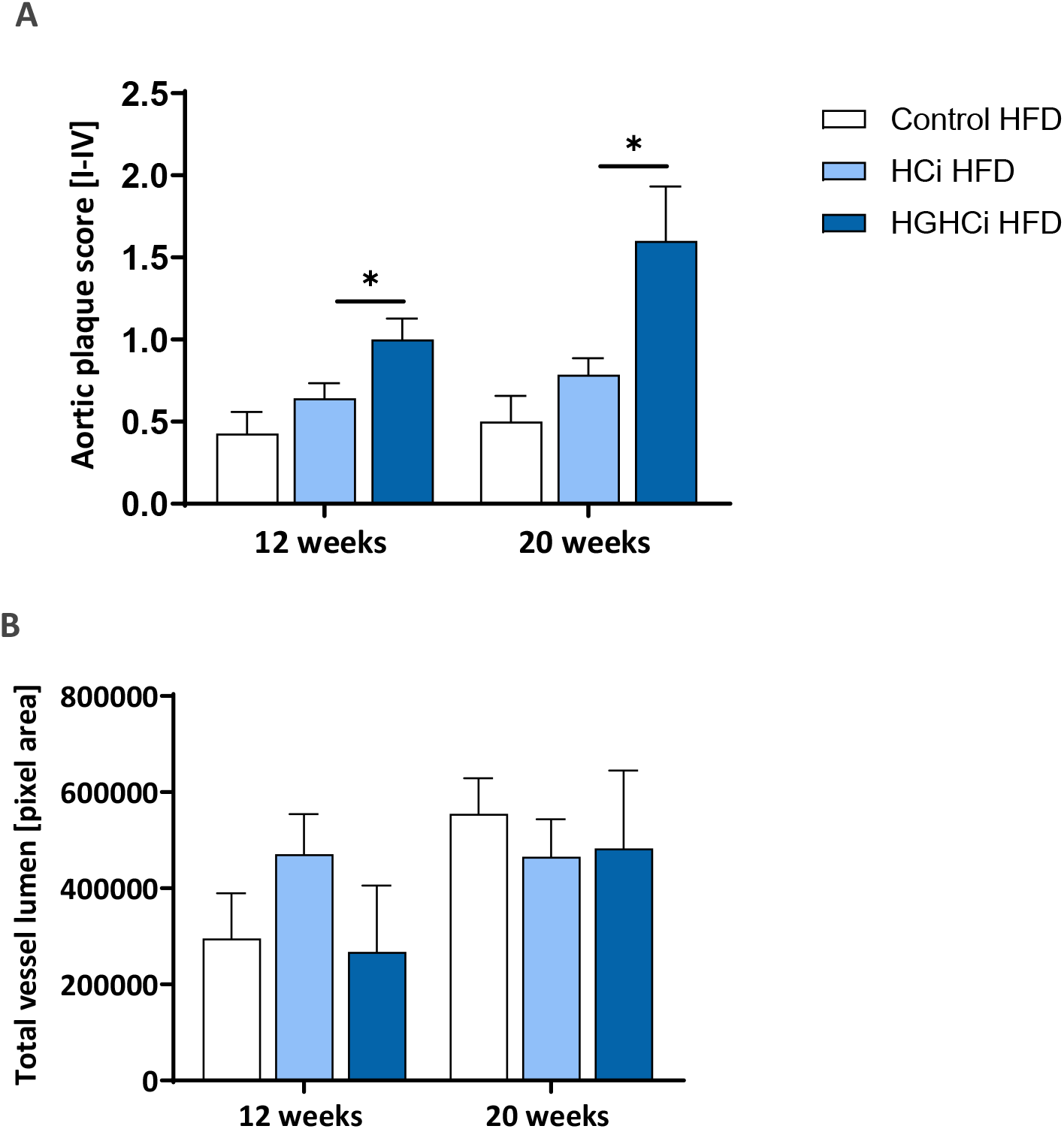
Comparison of aortic plaque score and total vessel lumen area. **A:** Aortic plaque score and total vessel lumen area [Pixel] of the truncus brachiocephalicus of HGHC and HCi mice on Western diet (high-fat diet, HFD) after 12 and 20 weeks. Aortic plaque score was determined as described below. 0= no lesions; 1 = Lesions only in bifurcation; 2= like 1 + at least one long-stretch lesion; 3= like 1 + at least two long-stretch lesions; 4= like 1 + three to four long-stretch lesions

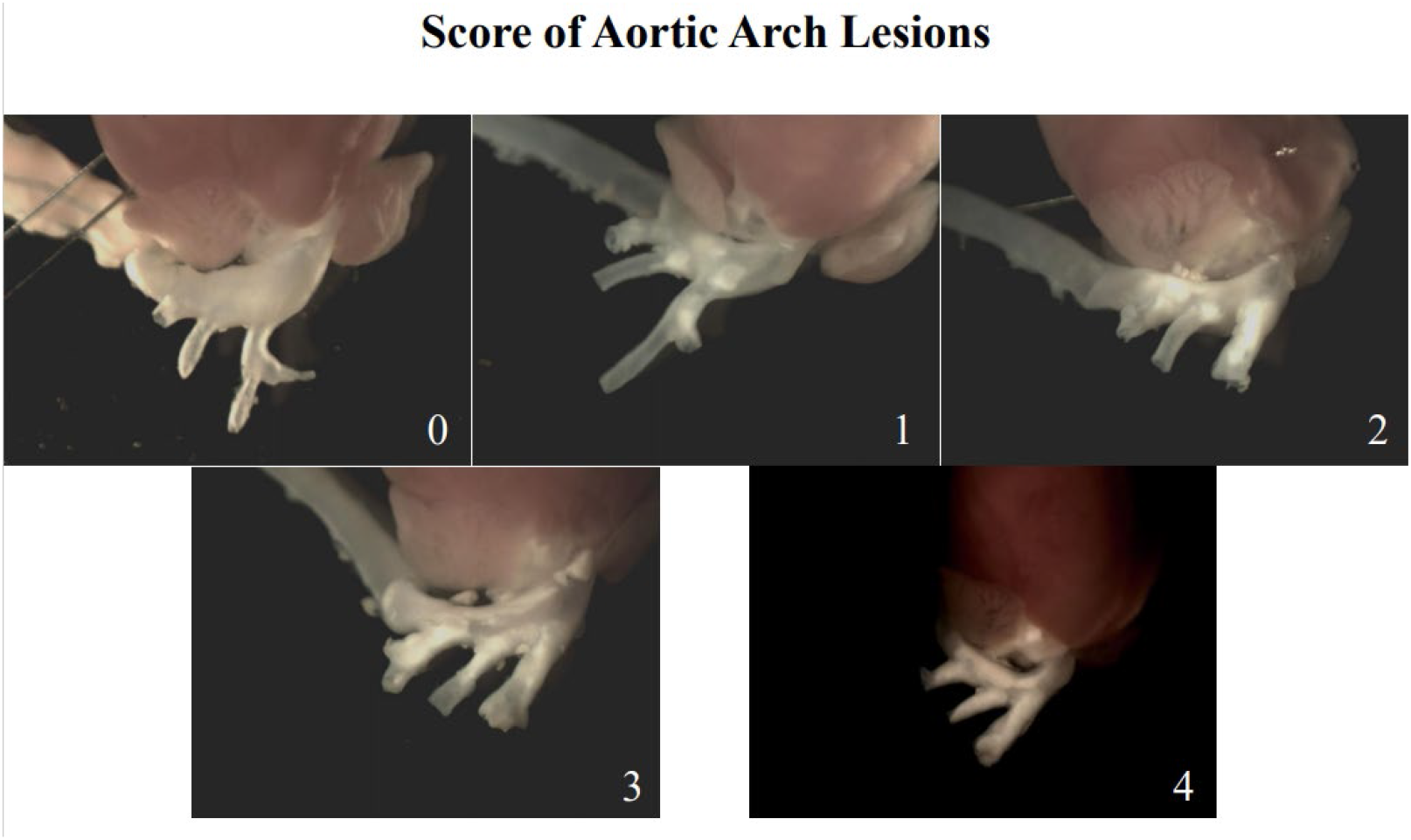

**Supplementary Figure IV.**
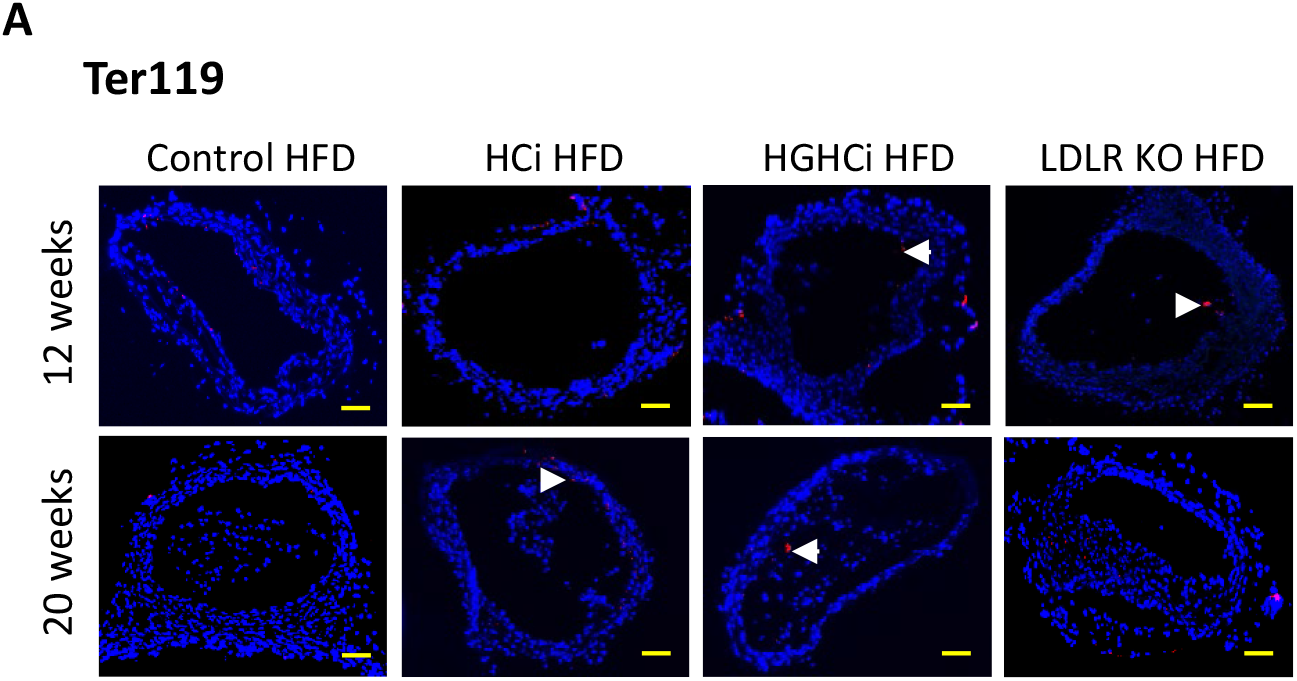
Intraplaque hemorrhage is not increased in HGHCi HFD mice. Representative images showing immunofluorescence staining of truncus brachiocephalic arteries sections for hemorrhage marker Terr-119 (**A,** Terr-119= red; DAPI nuclear counterstain= blue, white arrows). Scale bar 200 μm. Control 12 and 20 weeks (N= 7), HCi 12 weeks (N= 5) and 20 weeks (N= 6), HGHCi 12 weeks (N= 7) and 20 weeks (N= 6). Control HFD: Wild type mice without rAAV8-PCSK9^D377Y^ injection on high fat diet (HFD); HCi HFD: rAAV8-PCSK9^D377Y^ injection plus HFD (hyperlipidemic); HGHCi HFD: rAAV8-PCSK9^D377Y^ and streptozotocin injection and HFD (hyperlipidemic and hyperglycemic).

**Supplementary Figure V.**
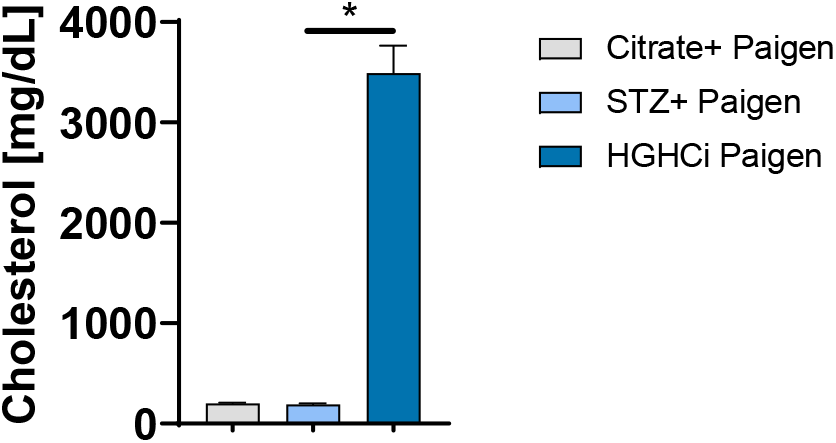
Hyperglycemia alone does not affect plasma cholesterol levels in mice fed a Paigen diet. Comparison of cholersterol levels in HGHCi mice (N=13), streptozotocin (STZ) treated mice (N=5) and citrate controls (N=3) fed with Paigen diet. Plasma cholesterol leves [mg/dL] were analyzed 12 weeks after induction of hypergylcemia with STZ.

